# Specific, replicable behavioral and neural correlates of sensory over-responsivity in childhood

**DOI:** 10.1101/2025.09.04.672422

**Authors:** Hexin Luo, Andrew W Kim, Christina A Gurnett, Anna M Abbacchi, John N Constantino, Joan L Luby, Michael T Perino, Deanna M Barch, Chad M Sylvester, Catalina M Camacho, Rebecca F Schwarzlose

## Abstract

**Objective:** Sensory over-responsivity (SOR), characterized by strong negative reactions to typically innocuous stimuli, is considered a symptom of autism spectrum disorder. However, SOR also affects 15-20% of children overall, including a majority of children with common psychiatric conditions. Despite its prevalence, the clinical specificity and neurobiological bases of SOR remain poorly understood. Our study aims to determine the specific clinical significance of SOR across diverse child samples and establish whether SOR is associated with replicable patterns of functional connectivity (FC).

**Method:** We analyzed data from 15,728 children (ages 6-17.9 years) across five datasets: three community samples, including the Adolescent Brain Cognitive Development [ABCD] and Healthy Brain Network [HBN] studies, and two autism-enriched samples. Bivariate and multivariate models examined associations between SOR and symptoms of anxiety, attention-deficit/hyperactivity disorder, depression, conduct disorder, and oppositional defiant disorder, as well as autistic traits. Analysis of resting-state functional MRI (fMRI) data from the ABCD study (n=4195) identified candidate brain-wide and circuit-specific FC correlates of mild SOR and tested replication of effects in an independent ABCD subsample (n=4190). Additional analyses tested extension to children with severe SOR in ABCD and replication in smaller samples [HBN (n=356), ABCD subset (n=356)].

**Results:** Multivariate analyses revealed that SOR is associated with a remarkably consistent profile across samples: greater levels of both autistic traits and anxiety symptoms and, in community samples, lower levels of conduct disorder symptoms. Across samples, SOR is not reliably associated with symptoms of any other analyzed psychiatric conditions. Mild SOR is associated with brain-wide FC patterns and, specifically, reduced FC between cingulo-parietal network and bilateral caudate nucleus; these patterns show robust replication across independent ABCD subsamples and extension to severe SOR in ABCD but are not significant in smaller samples, indicating that large sample size is needed to reliably detect brain effects.

**Conclusion:** Results suggest that SOR may constitute a latent trait associated with both specific clinical risk and protection, and with replicable cortico-subcortical neural correlates. These findings advance our understanding of the neurobiology and clinical relevance of SOR. They may also inform clinical practice and future research aimed at understanding and supporting individuals with sensory challenges.

## Introduction

Sensory over-responsivity (SOR) is a behavioral pattern characterized by strong, negative responses to sensory stimuli that are typically considered innocuous, such as traffic noise or tactile stimulation from clothing. SOR affects approximately 15 to 20% of children^1–4^ and is associated with psychiatric problems and significant functional impairment, including sleep and gastrointestinal problems.^5–8^ Studies linking SOR to enhanced physiological responses to sensory stimulation and altered activation of sensory brain areas have provided early clues that SOR may reflect differences in sensory processing and habituation.^3,9–15^ Parallel work from animal models shows that neonatal transgenic mice with autism-associated mutations exhibit peripheral and central sensory dysfunction that, in turn, triggers a developmental cascade to long-term anxiety-like behaviors and social deficits.^16–23^ Interventions to mitigate this dysfunction in early life rescue mice from anxiety-like behavior and reduce social impairments later in development, demonstrating a causal role for sensory dysfunction in long-term behavioral and social challenges.^24^ Because SOR tends to manifest in early childhood,^4,25,26^ before the onset of most psychiatric conditions, and is readily observed by caregivers, it may be a useful marker for identifying high-risk children who would benefit from psychiatric screening. However, its associations with diverse conditions raise an important question: is SOR a universal correlate of psychiatric conditions, or does it provide more specific information about clinical symptoms and risk?

Despite the prevalence of SOR in childhood and its relation to functional impairment, there remains no consensus across clinical specialties regarding whether or how SOR relates to clinical diagnoses. SOR has long been recognized for its strong association with autistic traits and was added as a symptom of autism spectrum disorder (ASD) in The Diagnostic and Statistical Manual of Mental Disorders, Fifth Edition (DSM-5) used for psychiatric diagnoses.^27,28^ However, SOR is also over-represented among children with neuropsychiatric conditions beyond autism, including anxiety disorders, attention-deficit/hyperactivity disorder (ADHD), tic disorders, and obsessive-compulsive disorder (OCD), and is understood to reflect risk for emerging psychopathology.^2,29–34^ SOR has also been conceptualized as a central feature of sensory modulation challenges, a type of regulatory-sensory processing disorder that is included in the Diagnostic Manual for Infancy and Early Childhood-Revised^35^ but not the International Classification of Diseases 10^th^ Revision (ICD-10)^36^ or the DSM-5.^37^ In short, there is no clinical and scientific consensus about whether SOR should be conceptualized as a symptom of a specific neurodevelopmental condition (i.e., ASD), a non-specific correlate of psychiatric and neurodevelopmental conditions, a defining feature of a standalone sensory processing disorder, or, as suggested by studies of animal models, a risk factor that promotes a specific constellation of symptoms in childhood. Determining the specificity of SOR associations with psychiatric symptoms across heterogeneous child samples can inform this important debate and advance our understanding of SOR.

Existing research linking SOR to psychiatric conditions has largely taken a bivariate analytic approach by examining the relation of SOR with symptom burden or diagnostic status for a single condition. However, interpretation of results from this research is limited by the considerable comorbidity and subclinical symptom associations of common conditions of childhood, such as autism, anxiety, and ADHD.^8,38–41^ Given this overlap, a multivariate approach is needed to determine whether SOR indicates universal versus specific psychiatric problems.

We previously applied this multivariate approach to data from a large, community sample of preadolescents and found significant SOR-symptom relationships in both positive (anxiety, depression, oppositional-defiant disorder) and negative (conduct disorder) directions.^3^ Yet due to the extreme size (N>11,000) of the sample, even small (i.e., clinically insignificant) effects can achieve statistical significance. Moreover, those effects may be specific to that sample’s clinical or demographic characteristics, including its restricted age range (10-11 years). A multi-sample approach is therefore needed to determine whether robust, specific SOR-symptom associations exist for conditions other than autism.

Like many behavioral phenotypes associated with psychiatric conditions, SOR is a dimensional trait that exhibits a continuum of severity and associated functional impairment across the population. Whereas severe SOR has received more clinical and research attention, mild SOR appears to be more prevalent in the general population, constituting approximately two-thirds of children with SOR.^3^ It remains an open question whether mild and severe SOR have shared clinical and neurobiological correlates, although they are generally believed to reflect the same underlying phenotype. Moreover, to date, there is no universally accepted cutoff to demarcate mild versus severe SOR. Our recent analysis of functional magnetic resonance imaging (fMRI) data from the Adolescent Brain Cognitive Development (ABCD) dataset provided insights into the neural correlates of severe SOR, including identifying differences in functional connectivity (FC) patterns in the brains of children with severe SOR, relative to children without SOR.^3,11,42^ These differences included reduced FC within somatosensory brain networks and increased FC between somatosensory and salience networks, highlighting candidate alterations in neural circuitry that may contribute to severe SOR. However, because fewer than 6% of children in the sample had severe SOR, we could not test replication of these findings in an independent subset of the sample. By contrast, mild SOR was reported in 12.4% of children in the sample, yielding sufficient numbers for conducting a data-driven, whole-brain analysis of FC-SOR associations with independent subsamples of children for analytic exploration and replication. We note that prior analyses did not examine FC data from children with mild SOR. In the present study, we ask whether FC relations with mild SOR are sufficiently robust to replicate in independent subset of the ABCD sample. To our knowledge, no data-driven, whole-brain FC analysis of SOR has been tested for replication in an independent subsample. If SOR is a dimensional trait, children with mild SOR might be expected to show similar neural effects as children with severe SOR, albeit potentially with smaller effect sizes. Alternately, FC patterns associated with severe and mild SOR may not the same. Therefore, the current study additionally tests whether FC correlates of mild SOR that replicate across subsamples also extend to severe SOR.

The current study analyzed 15,698 children across five samples that differ in behavioral, clinical, and demographic features to 1) determine the specific clinical symptom significance of SOR across pediatric samples; and 2) establish whether SOR is associated with distinct FC patterns that replicate across independent sets of children. To address the first aim, we conducted parallel analyses of SOR and psychiatric symptoms across multiple community (Human Connectome Project-Development [HCP-D]^43^; Healthy Brain Network [HBN]^44^; ABCD^45^) and autism-enriched (Simons Simplex Collection [SSC]^46^; Brain Gene Registry [BGR]^47^) samples and computed pooled effect sizes to identify effects that were robust and generalizable. To address the second aim, we capitalized on the large number of children ages 10-11 years with mild SOR in the ABCD dataset to identify candidate associations between FC and mild SOR in an exploratory subset of the sample (ABCD exploratory: n=554 mild SOR, n=3641 no SOR) and then test replication of these associations in an independent subset of ABCD participants (ABCD replication: n=457 mild SOR, n=3665 no SOR). Given that many brain-behavior associations exhibit small effect sizes^23^ and that psychiatric symptoms often result from many small alterations in FC patterns throughout the brain, we first examined whether mild SOR is associated with replicable brain-wide FC patterns across the exploratory and replication subsamples. To provide clues about circuit-specific differences, we identified specific SOR-FC associations in the exploratory subsample and tested replication of these associations in the independent subsample. Candidate FC-SOR associations identified in the exploratory subsample were also tested for extension to a substantially smaller dataset enriched for behavioral problems. Findings from this work provide valuable information about childhood SOR that could prove useful for conceptualizing the role of SOR in psychiatric and neurodevelopmental conditions, as well as for early risk identification and screening.

## Method

### Samples for Symptom Analyses

To identify SOR-symptom associations that are robust across sample characteristics, data were obtained from five independent datasets (total *N*=15,728; ages 6-17.9 years, Table 1) that vary dramatically in psychiatric symptoms, autistic traits, and rates of SOR. These datasets include community samples, clinical samples, and samples enriched for psychopathology. Specific sample sizes vary by analysis depending on data availability.

**Table 1:**
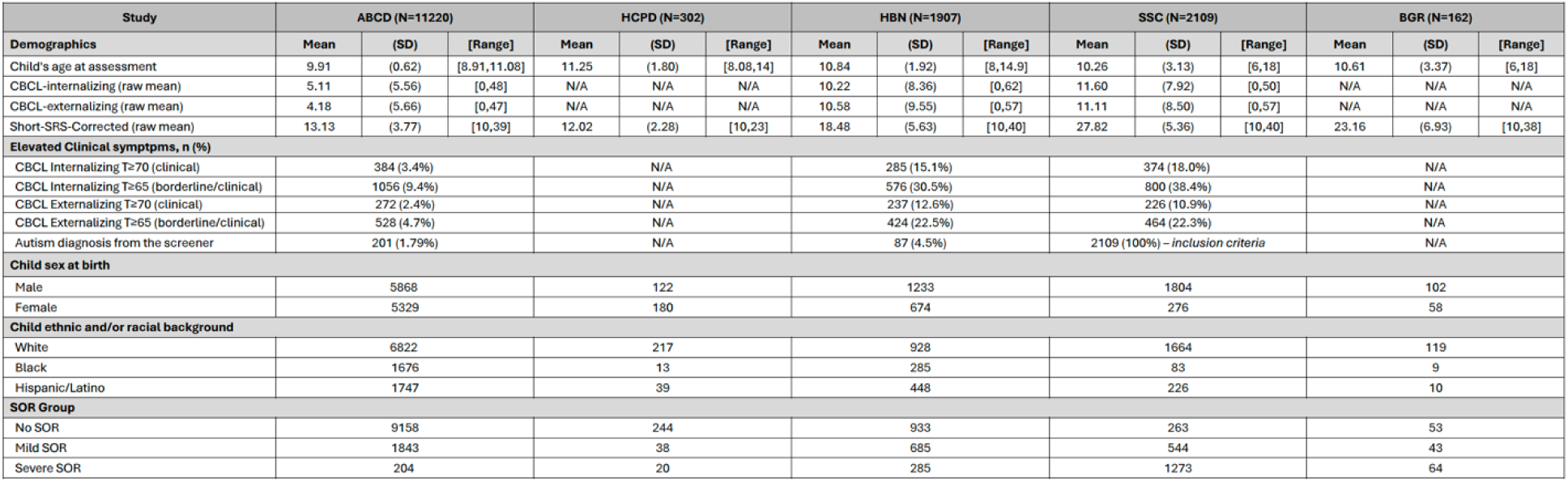
Summary of participant demographic and symptom information by dataset. N/A means data is not available at the time of writing.

#### The Human Connectome Project-Development Study (HCP-D)

HCP-D is a cross-sectional community study of children recruited to represent normative development across diverse demographic backgrounds while excluding individuals with serious medical conditions. ^43^ Given the breadth of ages in this study and the low numbers of children under the age of 8, analyses for the current study were restricted to *N*=302 participants ages 8-15 years with all required measures collected.

#### The Healthy Brain Network Study (HBN)

HBN is a cross-sectional, multi-site study enriched for children with behavioral problems. ^48^ The current study analyzed data from *N*=3,635 HBN participants ages 8-14 with all required measures collected.

#### The Simons Simplex Collection (SSC)

The SSC is a core project of the Simons Foundation Autism Research Initiative that collected data from families with a child affected by autism.^46^ Data from *N*=2,109 children ages 6-18.9 years with autism diagnoses and all required measures were included in the current analyses.

#### The Brain Gene Registry (BGR)

BGR is a national repository of information on individuals with variants in genes related to intellectual and developmental disabilities.^47^ Behavioral and symptom information is additionally collected and stored in the repository when possible. Data from the *N*=162 participants aged 6-18.9 years (mean: 10.6 years) with available measures were analyzed for the current study.

#### Adolescent Brain Cognitive Development ^SM^ Study (ABCD Study^®^)

ABCD study is a longitudinal, multi-site community study of adolescents recruited through a school-based strategy designed to maximize sample representativeness, beginning at ages 9-10 years.^45,49^ Analyses were restricted to children with all required measures from the baseline timepoint (*N*=11,218) from data release 5.1^50^ (with documented labeling errors ^51^ in tabulated rsfMRI network-subcortical regions of interest addressed). ABCD study recruitment neither excluded nor enriched based on psychiatric conditions or autism. However, children with conditions that required special schooling were not recruited. We reported results of related analyses from this dataset previously.^3^ Reanalysis was performed to provide consistency in modeling approach and permit pooling of effect-size estimates across datasets in the current study.

### Assessments

To maximize the interpretability of results across datasets, we restricted analyses to the same core set of standardized parent-report measures collected from all included samples. Additional details about these measures are provided in the Supplement.

#### Child Behavior Checklist (CBCL 6-18)

The CBCL 6-18 is a Likert-style, norm-referenced caregiver-completed rating scale that assesses children’s functioning (ages 6-18) during the previous six months.^52^ For each item, parents/guardians respond using a 3-point Likert scale: “Not True” (0), “Somewhat or Sometimes True” (1), and “Very True or Often True” (2). For modeling of SOR-symptom associations, we analyzed the raw scores from DSM-oriented subscales of the CBCL, which have demonstrated strong validity for screening and measuring specific psychiatric problems. ^52–55^ These subscales comprised depressive problems, anxiety problems, attention-deficit/hyperactivity problems, oppositional defiant problems, and conduct problems. Models for FC analyses based on the neuroimaging data included the broader internalizing and externalizing problem subscales, which represent higher-order dimensions of psychopathology.

#### Social Responsiveness Scale, Second Edition (SRS-2)

Parent report of child SOR was obtained from the SRS-2, a parent-report instrument used to identify the presence and severity of social impairment and autistic traits.^56^ Analyses were limited to the 11 items from the SRS-2 that were administered in all five datasets. As described previously,^3^ responses on one item from SRS-2 (*My child seems overly sensitive to sounds, textures, or smells)* were used to assign each participant to a SOR category. Item responses were given on a four-point scale: “Not True” (0), “Sometimes True” (1), “Often True” (2), or “Almost Always True” (3). Given known limitations for modeling ordinal outcome variables,^57^ the single-item SOR measure was handled categorically for subsequent analyses; specifically, children were categorized into three SOR groups: No SOR (0), Mild SOR (1), and Severe SOR (2 or 3). These groups demonstrate a clear stepwise pattern of increasing psychiatric symptom burden across the No, Mild, and Severe groups (See Table S1, Table S4, and Supplement text). In two other child samples with both the SRS-2 and a more extensive questionnaire assessing sensory behaviors (Sensory Experience Questionnaire, SEQ 3.0^58^), we confirm there is a strong positive association between the single item SOR measure and a multi-item composite SOR score (rho_1_(106)=0.55, rho_2_(78)=0.70, both p<0.0001; see Supplement) and stepwise increases in multi-item SOR scores across SRS-derived No, Mild, and Severe SOR groups (see Supplement), indicating that the single-item measure has convergent validity with an established, multi-item measure of SOR. A dimensional measure of autism traits excluding SOR was derived from the sum of the remaining ten SRS-2 items that were administered across all five studies. The summed raw scores were converted to standardized T-scores for cross-sample comparison and statistical analysis.

### Participants for Neuroimaging Analyses

Primary functional MRI analyses were conducted on data from the ABCD Study, given the large number of participants with mild (*n*=1,069) or no SOR (*n*=7,306) in this sample, as defined above. To meet best practices of replication and reproducibility, we generated independent matched exploratory and replication subgroups from ABCD participants with no or mild SOR through random assignment followed by iterative case-reassignment until the subgroups were matched on demographic, psychiatric, and SOR characteristics (see Tables S1, S2 for full details). After excluding participants missing functional data, we retained *N*=4,195 for the exploratory group and *N*=4,190 for the replication group. Each group was comprised of approximately 13% participants with mild SOR (*n*=554 and *n*=525, respectively) and the remaining 87% of participants with no SOR. We additionally tested whether FC-SOR effects found for mild vs no SOR in the exploratory sample extended to *n*=457 children with severe SOR. Follow up analyses tested whether the pairwise FC-SOR results from ABCD could be detected in substantially smaller samples using both a small, separate study with distinct characteristics (i.e., sample age range, sample psychopathology, sample size, and task during data collection) from HBN participants 8-11 years of age (*N*=356) with high-quality fMRI data and SOR measures,and a subset of ABCD participants (N=326) matched to the HBN sample in sample size and clinical profile (see Supplement for full details). Finally, to ensure that results were not due to family-level clustering across subsamples, analyses were repeated with the ABCD reproducible matched samples (ARMS) groups^59^ that maintained strict separation of families between subsamples (see Supplement).

### Neuroimaging Data Acquisition and Preprocessing

Resting-state functional MRI (rs-fMRI) data were collected using 3T scanners across 21 ABCD sites at the baseline timepoint, adhering to detailed protocols published previously.^45^ Children underwent high-resolution (2.4-mm isotropic, repetition time = 800 ms) multiband (factor = 6) scanning during rs-fMRI acquisition. Scan sessions typically included three or four 5-minute runs of rs-fMRI, with slices acquired in the axial plane.

Standardized preprocessing of rs-fMRI data was performed, encompassing registration, distortion correction, and normalization. Post-processing for resting-state FC involved several steps: regression of 24 temporally filtered motion parameters, exclusion of outlier frames with framewise displacement >0.2 mm, and regression of signals from white matter, cerebrospinal fluid, and the whole brain. Cortical parcels were delineated and assigned to one of 13 functional networks according to the Gordon–Laumann parcellation scheme,^60^ whereas subcortical regions of interest were identified using the ASEG FreeSurfer subcortical segmentation scheme.^61^ Mean pairwise blood oxygen level–dependent (BOLD) signal correlations between cortical parcels, as well as between cortical parcels and subcortical regions, were computed, Fisher-transformed, and averaged by cortical network (422 total FC pairs: 103 cortical-cortical network pairs and 319 cortical-hemisphere-specific subcortical pairs). Analyses were limited to data from participants recommended for inclusion based on ABCD study’s recommendations for data quality control.^62^ See supplement for additional details about neuroimaging data acquisition and preprocessing in both ABCD and HBN.

### Main Analysis

#### SOR-Symptom Analyses

We examined the relationship between SOR and psychiatric symptoms across five datasets, encompassing 15,728 participants ages 6-18 years. For large, cross-sectional datasets (HBN and SSC) with broader age ranges, we further subdivided participants into specific age groups (see Table S3 for details of subgroups) to allow for developmental variations, resulting in 9 separate subsamples of participants. For each subsample, we conducted separate hierarchical mixed-effects models comparing no SOR with SOR (either mild or severe). SOR was analyzed as a categorical rather than ordinal variable due to the known poor fit of ordinal predictors in hierarchical mixed-effects modeling.^57^ This categorical specification optimizes model convergence and interpretation while avoiding the statistical complications inherent in ordinal MLM specification. We separately computed effect-size estimates from conventional bivariate approaches and from our multivariate approach that accounts for common comorbidity/symptom co-occurrence. For bivariate analyses, we tested each psychiatric symptom individually (anxiety, depression, ADHD, ODD, conduct disorder, and autism). In the multivariate analysis, we tested all six psychiatric symptoms together in a single mixed-effects linear model with SOR. In all models, the dependent variable was binary SOR group, with psychiatric symptoms scores entered as continuous predictors. For datasets that included site information (ABCD, HCP-D, and HBN), we incorporated data collection site as a second-order random-effects variable in the models to account for site-specific variations in our analyses.

Given that SOR is considered a symptom of autism, its clinical significance might differ for autistic and community samples. We therefore grouped studies into community (ABCD, HCP-D, HBN) or autism (SSC, BGR) datasets based on their study recruitment objectives and severity of autistic traits in the sample. Odds ratios from bivariate and multivariate models were pooled separately across community and autism samples to assess the overall strength and consistency of SOR-symptom associations within these distinct populations. To confirm that the pattern of results was not contingent on SOR grouping, we also performed the analyses separately for mild versus no SOR and for severe versus no SOR (see Supplement for details). Analyses were carried out using the *nlme* and meta packages available in R (version 4.3.2).^63^

#### Functional Connectivity Analyses

We adopted a two-stage approach to FC analyses. First, we analyzed FC patterns across the whole brain to determine whether these global patterns are associated with mild SOR. We then tested whether specific FC pairs demonstrate robust, replicable associations with mild SOR, providing insight into circuit-specific differences. For both analytic approaches, we used independent subsamples to identify and replicate FC-SOR associations and additionally tested their extension to severe SOR.

We evaluated associations between mild SOR and FC in the exploratory, replication, and severe subgroups of the ABCD dataset using hierarchical linear mixed-effects models of FC that accounted for scanner-related variance across multiple study sites as a second-level random-effects variable. We tested for associations with SOR by modeling pairwise (network-network or network-subcortical structure) FC for all FC pairs (see Supplement for a complete list of regions tested). In addition to the second-order variable of scanner, the model contained key covariates, including sex, age, economic disadvantage, framewise displacement, psychiatric symptoms scales (standardized CBCL-Internalizing, CBCL-Externalizing raw scores) and autistic traits (adjusted S-SRS score excluding SOR item) as first-order variables. This comprehensive approach allowed us to disentangle the specific contribution of SOR from related demographic, clinical, and methodological factors, thereby isolating its unique effect on functional connectivity while controlling for potential confounding variables. Tabulated resting state functional connectivity, SOR, and all covariates were mean-centered and transformed into standardized scores, and then winsorized using the interquartile method.

#### Brain-wide Functional Connectivity Pattern Analysis

From each dataset, we extracted standardized beta coefficients from multilevel models (MLMs) representing the magnitude and direction of the relationship between SOR and FC pairs. These coefficients were organized into symmetric matrices where both rows and columns represented the 32 brain regions of interest (13 cortical networks and 19 subcortical structures). Each cell (*i,j*) in the matrix contained the beta coefficient reflecting the SOR-related FC between regions *i* and *j*. To quantify the similarity of SOR-related FC patterns across exploratory and replication subsamples, we calculated Spearman rank correlations (ρ) between the beta coefficient matrices. Two key comparisons were performed: ABCD Exploratory vs. ABCD Replication and ABCD Exploratory vs. ABCD Severe.

To assess the statistical significance of cross-sample beta correlations, we implemented a permutation testing approach by creating 1,000 permuted versions of the exploratory ABCD subsample in which the SOR variable was randomly shuffled while preserving the original distribution. For each permutation, we generated new beta coefficient matrices using the same MLM model, then calculated Spearman correlations between these permuted matrices and the original matrices from comparison datasets (ABCD Replication, ABCD Severe). The *p*-value for each cross-dataset comparison was calculated as the proportion of permuted correlations that were greater than or equal to the actual observed correlation.

#### Identifying Specific Candidate FC Pairs Associated with SOR

To identify robust, SOR-specific candidate brain regions in the exploratory subsample, we applied a MLM to each pairwise FC variable using a primary SOR+Demographics model that included SOR, demographic variables (sex, age, economic disadvantage), and mean framewise displacement (FD) to control for motion effects. To ensure that findings are robust to model selection, we additionally computed basic models (Minimal Model: SOR only) and models with additional psychiatric covariates (Symptom Model: SOR+Demographics+FD+Symptoms model that included internalizing/externalizing behaviors and autistic traits; see Supplement). P-values for fixed effects in each model in the exploratory sample were adjusted for multiple comparison using the Benjamini-Hochberg procedure, with significance determined using an adjusted alpha threshold of 0.01. Identifying a given candidate FC pair in the exploratory subsample or determining that it replicated in the replication subsample required statistical significance (p<.01) of SOR effect in both the minimal model and the primary SOR+Demographics model.

#### Testing Extension of Results to Severe SOR in ABCD

Although prior work investigated FC associations with severe SOR in ABCD, those analyses did not test effects from all pairwise FC variables or assess replication in a sample of independent participants.^3^ The current investigation of FC associations with mild SOR included exhaustive pairwise exploratory analyses and independent replication. To determine if FC associations with mild SOR identified in the current exploratory analyses extend to severe SOR, we therefore applied the same MLM models to a subset of the ABCD sample comprising children with severe SOR and the participants with no SOR from the replication subsample.

## Results

### SOR and psychiatric symptoms by dataset

SOR rates varied substantially across samples (Figure 1A). Rates of SOR (combined mild and severe) in community samples were 19.2% in the HCP-D dataset, 18.3% in the ABCD dataset, and 51.0% in the HBN dataset. In samples enriched for autism, rates were 87.3% in the SSC dataset and 66.9% in the BGR dataset. Profiles of dimensional clinical symptoms also varied dramatically across samples (Figure S1). As expected, we observed positive associations among all dimensional scores of psychiatric symptoms (anxiety, depression, conduct disorder [CD], attention-deficit/hyperactivity disorder [ADHD], oppositional defiant disorder [ODD], and autism spectrum disorder [ASD]) within samples (See Figure 1B, Figure S2).

**Figure 1:**
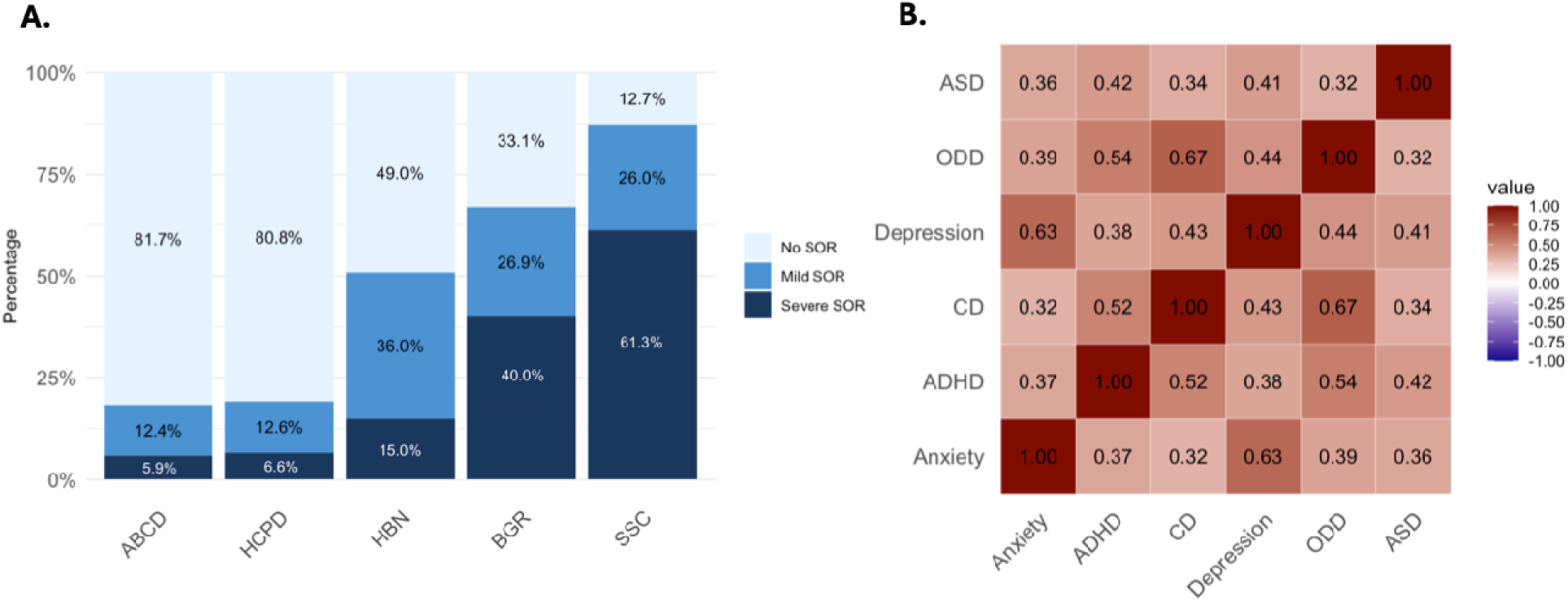
SOR and psychiatric symptoms measures in five independent datasets. A. Rates of SOR group by dataset. B. Matrix of correlations computed between clinical symptom scores within datasets and then averaged across datasets.

### SOR-Symptom Associations

Results from conventional bivariate analyses show indiscriminate positive associations between SOR and all psychiatric measures (all pooled ORs >1.2, *p* < 0.05; Figures S3 and S4). However, the multivariate models that simultaneously incorporate all symptom variables revealed a striking pattern: specific, independent associations of both anxiety symptoms and autistic traits with SOR. Notably, anxiety is significantly associated SOR (pooled OR = 1.50, 95% CI [1.42, 1.58]; Figure 2) with mild heterogeneity (I²= 33.2%) in effect pooled across all subsamples.

**Figure 2.**
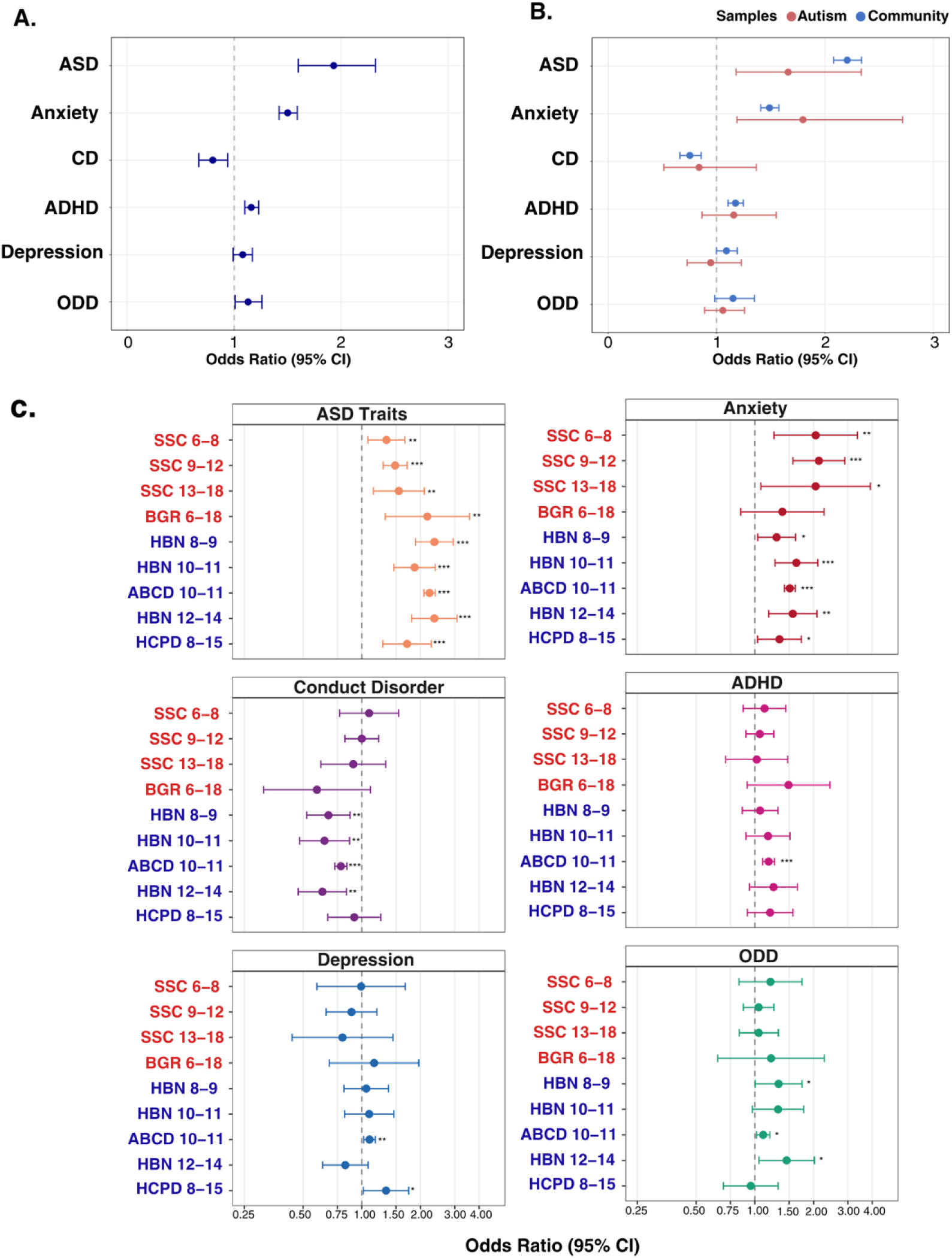
SOR is consistently associated with both anxiety and autism. A. Multivariate associations between SOR and psychiatric symptoms pooled across all subsamples. B. Multivariate associations between SOR and psychiatric symptoms by community and ASD samples. C. Age-specific multivariate associations of SOR with all modeled symptoms. Community samples are indicated in blue font, autism samples in red font. Labels include subsample age ranges (in years) [**p < 0.01, ***p < 0.001.].

Results are similar when pooled by sample type (community samples: pooled OR = 1.48, 95% CI [1.40, 1.57], I² = 0%, and autism samples: pooled OR=1.93, 95% CI: [1.56, 2.40], I² = 1.7%), indicating highly stable effect sizes across different studies and age groups (Table 2). The confidence intervals for pooled effect size estimates of anxiety-SOR associations by SOR severity overlapped in community and autism samples, indicating no significant difference in the strength of anxiety-SOR associations across these two types of samples. We note that odds ratios from these multivariate models reflect the size of effects in terms of unshared variance and will therefore be smaller than odds ratios from bivariate models.

**Table 2.** Pooled effect sizes for associations of psychiatric symptoms with SOR in multivariate models computed across all nine subsamples. Confidence intervals shown in square brackets. I^2^ indicates heterogeneity. Significant associations (where the 95% CI does not include 1.0) are indicated [**p < 0.01, ***p < 0.001]. Community samples include HBN, ABCD, and HCP-D studies. Autism samples include SSC and BGR studies.

| **Symptom \ SOR** | **Pooled OR** | **I<sup>2</sup>** |
| --- | --- | --- |
| **ASD** |  |  |
| Community Sample | 2.17 [2.00; 2.35]*** | 28.3%, p=0.23 |
| ASD Sample | 1.48 [1.32; 1.65] *** | 5.4%, p=0.36 |
| **Anxiety** |  |  |
| Community Sample | 1.49 [1.41; 1.58]*** | 0%, p=0.57 |
| ASD Sample | 1.84 [1.56; 2.41]*** | 0%, p=0.52 |
| **Conduct Disorder** |  |  |
| Community Sample | 0.76 [0.67; 0.82]*** | 0%, p=0.25 |
| ASD Sample | 0.97 [0.83, 1.13] | 1.7%, p=0.38 |
| **ADHD** |  |  |
| Community Sample | 1.17 [1.10, 1.25]*** | 0%, p=0.90 |
| ASD Sample | 1.09 [0.96, 1.24] | 0%, p=0.61 |
| **Depression** |  |  |
| Community Sample | 1.08 [0.98; 1.19] | 37.7%, p=0.17 |
| ASD Sample | 1.07 [0.94, 1.22] | 0%, p=0.89 |
| **ODD** |  |  |
| Community Sample | 1.17 [1.04; 1.32]** | 0%, p=0.21 |
| ASD Sample | 1.07 [0.94; 1.22] | 0%, p=0.89 |

Multivariate analyses also supported the expected positive relation of SOR with other autistic traits (pooled OR across all subsamples = 1.92, 95% CI [1.60, 2.31])), but with high heterogeneity (I^2^= 86.1%). Intriguingly, the association between SOR and autistic traits was larger in the community samples (pooled OR=2.17, 95% CI: [2.00, 2.35], I² = 28.3%) than in the autism samples (pooled OR=1.48, 95% CI: [1.32, 1.65], I² = 5.4%). The multivariate analyses also replicated and extended published findings from the ABCD dataset^3^ that SOR is negatively associated with conduct disorder symptoms in models that account for co-occurring psychiatric symptoms (pooled OR across all subsamples= 0.79, 95% CI [0.67, 0.94], I² = 65.8%; Figure 2A). Separate pooling by sample type indicates that this association is specific to community samples (pooled OR = 0.74, 95% CI [0.67, 0.82], I² = 25%), whereas CD symptoms are not significantly associated with SOR in pooled analyses from autism samples (see Table 1 for full results).

Separate post hoc analyses for mild SOR and severe SOR (relative to no SOR) yielded a highly similar pattern of results (Figures S5 and S6). Any significant associations between SOR and psychiatric symptom other than anxiety or autism in the community samples were of small effect size. No other symptom scales beyond anxiety and autism were significantly associated with SOR in the autism samples. Supplemental multivariate models including sex and age as fixed effects produced the same results across all samples. Age was not a significant predictor of SOR for any sample, whereas male sex was associated SOR in the ABCD and HCP-D samples. See Supplement for additional details. This pattern of results demonstrates a *specific relationship between SOR and anxiety symptoms* that is consistent across nine subsamples varying in age, autism diagnostic status, and enrichment for other psychiatric symptoms.

### SOR and functional connectivity

The second aim of the current study is to advance neurobiological models of SOR by identifying FC correlates of SOR that replicate across independent sets of children. We undertook this test by focusing on mild SOR, which is sufficiently prevalent in the ABCD sample to support independent exploratory and replication subsamples of children with mild SOR. Multilevel models were used to derive beta weights for associations of brain-wide, pairwise network-level FC with mild SOR in an exploratory subset of children in the ABCD study and, separately, in a matched, non-overlapping replication subset of children with mild SOR in the study. The pattern of brain-wide FC-SOR betas replicated across these independent samples, as indicated by permutation testing (ρ=0.31, p_perm_=0.021; brain-wide FC-SOR effect: ρ=0.31, p<0.001; see Figure A,B and Figure S7). Analyses limited to the largest FC-SOR effects (top 10% of the absolute value of beta coefficients) in the exploratory sample yielded a higher correlation with the replication sample (ρ=0.61, p<0.001, Figure S8). FC-SOR effects from the exploratory sample for mild SOR also extended to severe SOR in an independent subset of children (ρ=0.55, p_perm_<0.001; brain-wide FC-SOR effect: ρ =0.55, p<0.001; top 10%: ρ =0.65, p<0.001), indicating that brain-wide patterns of FC effects may be shared across levels of SOR severity (mild vs severe).

Next, we utilized the exploratory and replication samples from the ABCD dataset to identify specific FC pairs that demonstrate replicable associations with mild SOR. Identification of such pairs can inform circuit-specific understanding of neural basis of SOR. Among the 422 FC pairs tested in the exploratory ABCD sample, nine met criteria for selection to be tested for replication in the replication sample (Figure 3C; Table S5). Of these FC pairs, two met the stringent criteria for replication (see Methods): mild SOR was associated with reduced FC between cingulo-parietal network and the left and right caudate nucleus (Figure 3D). Of the nine FC pairs identified in the exploratory sample, five met criteria for extension to the severe SOR subsample (Figure 3D; Table S5); these included the cingulo-parietal to bilateral caudate nucleus pairs mentioned above. Supplementary analyses with the ARMS subsamples confirmed that the replicating FC-SOR effects cannot be explained based on family-level effects (see Supplement). However, the effect sizes are small, and results appear to depend on large sample sizes, as we do not observe significant FC-SOR effects in the HBN (n=351, mean age: 9.82 years) sample, or in an ABCD subsample matched to the HBN sample (n=351, mean age: 9.45 years; see Supplement).

**Figure 3.**
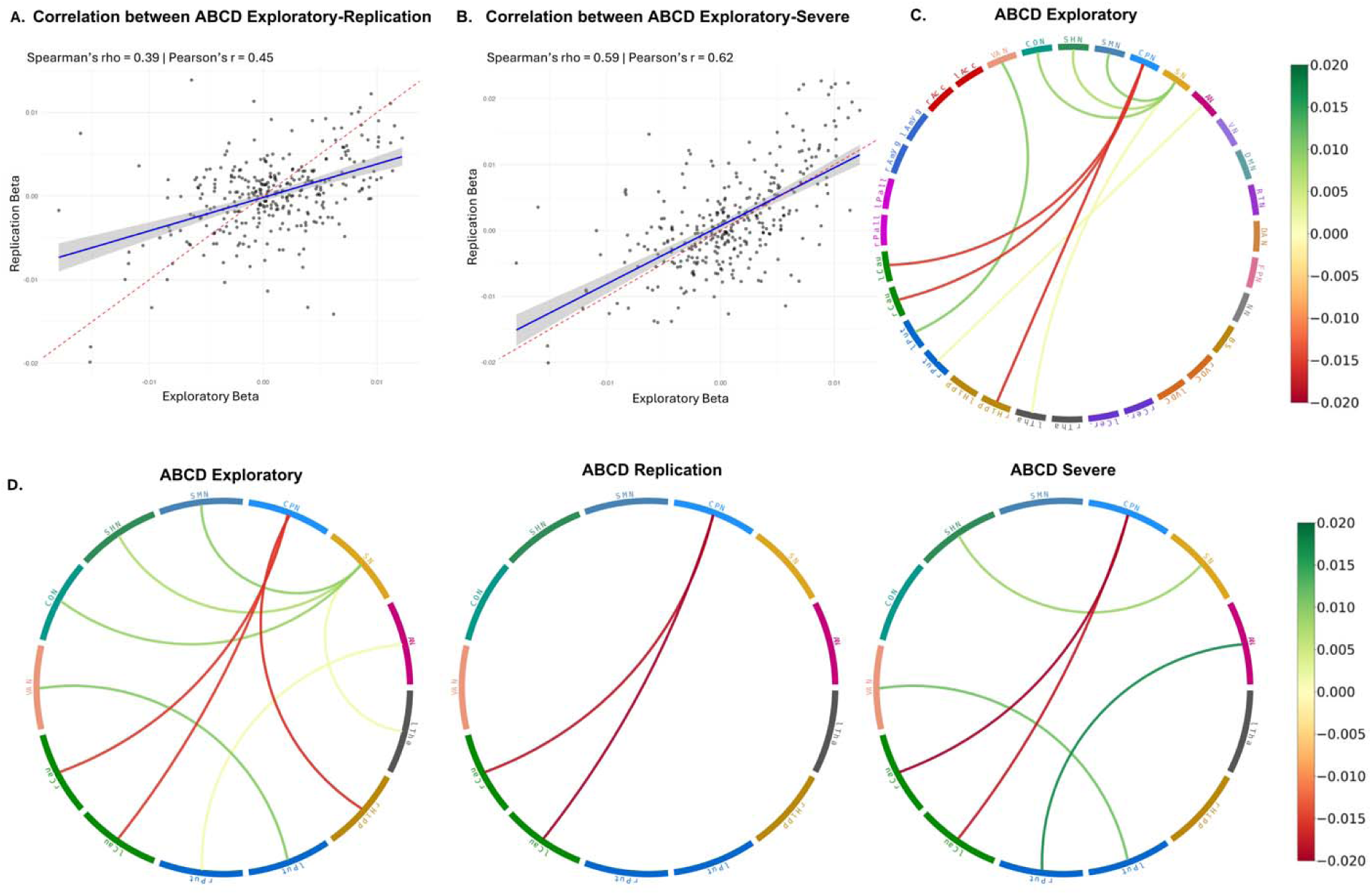
Brain-wide and specific, replicable FC differences patterns of FC-SOR associations across independent subsamples. A. Spearman correlations of functional connectivity pattern associated with SOR between ABCD exploratory-ABCD replication subsets. B. Spearman correlations of functional connectivity pattern associated with SOR between ABCD exploratory-ABCD Severe subsets. C. Change in functional connectivity associated with mild SOR from ABCD-exploratory SOR-demographics model. D. Replication test for functional connectivity pairs associated with SOR: Mild SOR (ABCD-Exploratory and Replication) and Severe SOR (ABCD-Severe) results from the SOR-demographics model

## Discussion

Based on data from more than 15,728 children across five independent datasets, this study provides new insights into clinical associations with common forms of SOR in childhood, demonstrating specific, consistent links between SOR, ASD and anxiety that transcend study design and population characteristics. Functional neuroimaging analyses of data from 9,197 children identified and independently replicated whole-brain and specific FC differences associated with SOR, establishing neural correlates and implicating new candidate neural circuits in the neurobiology of childhood SOR.

By clarifying the specific clinical relevance of SOR, the current findings inform our understanding of SOR and raise new questions. Given that SOR is neither universally associated with psychopathology nor uniquely associated with a single diagnosis, we might conceptualize SOR as a latent trait associated with higher likelihood for both autism and anxiety. A key outstanding question is exactly how SOR relates to these conditions. Are they causal relationships, and if so, what is the directionality of these effects? A few early findings suggest that SOR and associated sensory gating impairments may precede and promote emerging anxiety symptoms in development. These include correlational evidence from longitudinal studies showing that SOR unidirectionally predicts anxiety in children^2,9,10,64,65^ and experimental evidence that sensory gating disruptions in early development promote social impairments and anxiety-like behavior in transgenic mouse models of autism.^18,21,24,66^ Additional studies in humans and animal models are needed to resolve the nature, causality, and directionality of relationships between SOR and both autism and anxiety.

The present analysis of clinical symptoms demonstrates the importance of accounting for co-morbidities and co-occurring symptoms in pediatric psychiatry. Whereas bivariate analyses yield non-specific positive associations between SOR and symptoms of all psychiatric conditions analyzed, multivariate analyses reveal robust, specific associations between SOR and anxiety symptoms in both autistic and community samples. Multivariate models also uphold expected positive associations between SOR and autistic traits in most samples and support prior findings that conduct disorder symptoms are *negatively* associated with SOR in community samples when accounting for co-occurring symptoms.^3^ Although prior studies have documented higher rates of SOR among children with ADHD,^29,67,68^ the current results suggest that such findings primarily stem from the high comorbidity and symptom co-occurrence of ADHD with both anxiety disorders and autism.^39–41,68–71^

Results of the comprehensive exploratory neuroimaging analyses indicate that SOR is associated with brain-wide differences in functional neuroarchitecture and specific network-level differences. The analyses identified two robust FC-SOR effects that replicate across independent subsamples and generalize across SOR severity; negative associations of SOR with FC between the cingulo-parietal network and both the left and right caudate nucleus. The caudate nucleus, a subcortical nucleus of the striatum, participates in cortico-striato-thalamocortical circuits that support action initiation and diverse sensory and motor functions.^72^ The cingulo-parietal network, also known as the parietal memory network, is a relatively small cortical network encompassing the medial cingulate and posterior parietal regions that is thought to support memory processes.^60,73,74^ Based on the network’s FC profile in precision functional neuroimaging data, a recent study found that the cingulo-parietal network may unify with the salience network in individual participants to form a broader salience-related network.^75^ Although prior studies have focused on FC differences of the salience network in children with autism and in relation to SOR,^42,76–78^ the current findings specifically implicate FC differences in the cingulo-parietal network, opening up new avenues for future research.

Strengths of the current work include its large total sample size, analysis of effect consistency across several datasets differing in demographic and clinical sample characteristics, a data-driven, whole-brain investigation of FC correlates of SOR, and replication of identified FC effects in independent data. An important caveat to these findings is that analyses relied on basic parent-report measures of behavior and symptoms, including a single-item measure of SOR. Most large datasets do not include multi-item surveys of atypical sensory behaviors. Those that do employ a variety of instruments to measure sensory behaviors, making pooling across studies problematic. To harmonize cross-sample comparisons and support pooling of effect size estimates across samples, we used the SOR item present across these samples. However, this item only asks about SOR to sounds, textures, and smells, and therefore does not capture all forms of SOR. Future work in samples with more extensive measures of atypical sensory behavior are needed to determine whether the current findings replicate with these measures and extend to children with different (e.g., gustatory or visual) SOR triggers. In addition, we note that the no-SOR control group used in testing FC-SOR effects in the present analysis overlaps with the sample used in our prior publication examining FC pattern associate with severe SOR. Future work should seek to replicate these functional FC patterns across both SOR severity levels and control comparisons in entirely independent, large samples to further establish the robustness of these effects. Furthermore, while the current study demonstrates specificity in SOR associations with anxiety and ASD, future research should examine its relationship with other clinical conditions that have not been tested in the present study. For example, obsessive-compulsive disorder (OCD) and avoidant/restrictive food intake disorder (ARFID) both involve heightened sensitivity to sensory stimuli and may share overlapping profiles with SOR.^34,79–82^ The extent to which our findings generalize to or are distinct from these populations remains an open question.

We also note that although the FC-SOR associations identified in this study replicated in an independent dataset, effect sizes were small, and most available datasets would be underpowered to reliably detect them. Effect sizes are affected by a variety of factors, including the reliability of behavioral measures and functional connectivity metrics, which in turn vary based on the quantity of functional data acquired.^83^ They may also be affected by any heterogeneity of underlying neural signatures. Specifically, if heterogeneous neurobiological effects converge in affecting adaptive sensory modulation and producing the behavioral phenotype we know as SOR, then any specific neural difference would account for only a fraction of the cases of SOR, resulting in small overall effect sizes. Similar considerations are believed to affect FC associations with psychiatric symptoms more broadly.^84,85^ In light of this, replication of individual FC associations in the current study are noteworthy and specify a potential subset of the neural correlates of SOR, which can inform future research.

Our study provides evidence for the clinical and neurobiological significance of SOR in diverse populations through comprehensive analyses and replication across multiple datasets. The consistency in both clinical and neural signals suggests that SOR represents a unique, clinically relevant phenotype with distinct neural correlates. These insights can inform both clinical practice and future research aimed at understanding and supporting individuals with sensory processing differences.

