## Supplement for "Specific, replicable behavioral and neural correlates of sensory over-responsivity in childhood"

**Social responsiveness scale Second Edition (SRS-2) item-42 measuring SOR**

Our study used item 42 from the SRS-2-parent report: “My child seems overly sensitive to sounds, textures, or smells, to categorize SOR severity levels.” To validate this single-item measure, we examined its convergent validity in two different samples (BGR, n=108, Sensory Processing in Childhood Study (SPC: ages 9-14 years, N=80, unpublished data) which included both SRS-2 and a dedicated sensory assessment, the Sensory Experience Questionnaire (SEQ 3.0)-Parent report.^1^ In both the BGR and SPC samples, the SRS-derived SOR score showed a strong positive correlation with total SEQ sensory hyperreactivity score computed across sensory domains (rho(106)=0.55 and rho(78)=0.70, respectively; both p<0.0001) as well as composite sensitivity scores across the tactile, auditory, and olfactory modalities which the single-item question addressed (rho(106)=0.57 and rho(78)=0.72, respectively; ps<0.0001). We additionally conducted one-way ANOVAs and pairwise Welch's *t*-tests comparing SEQ 3.0 Sensory Hyperreactivity composite scores across the No, Mild, and Severe SOR groups derived from SRS-2 item 42 in two samples that contained both measures. In both samples, the one-way ANOVA was significant, [SPC: *F*(2, 77) = 32.78, *p* < .001, η² = .46; BGR: *F*(2, 104) = 24.38, *p* < .001, η² = .32]. For both samples, pairwise comparisons demonstrated that the No SOR group had significantly lower SEQ Hyperreactivity scores than both the Mild SOR group [SPC: *t* = −5.39, *p* < .001, *d* = −1.47; BGR: *t* = −4.84, *p* < .001, *d* = −1.21] and Severe SOR group [SPC: *t* = −7.44, *p* < .001, *d* = −2.02; BGR: *t* = −6.89, *p* < .001, *d* = −1.54]. Moreover, in both samples, the Mild SOR group had significantly lower SEQ Hyperreactivity scores compared to the Severe SOR group [SPC: *t* = −2.21, *p* = .032, *d* = −0.66; BGR: *t* = −2.60, *p* = .012, *d* = −0.60]. These results indicate that the single-item measure obtained from the SRS-2 has good convergent validity with a multi-item composite measure of SOR across sensory modalities.

We additionally assessed if the SOR severity groups (i.e., none, mild, and severe) derived from the single-item measure show significant differences in clinical symptom burden. Given the non-normality of score distributions, pairwise Wilcoxon rank sum tests were used to compare CBCL internalizing and externalizing T-scores across the No SOR, Mild SOR, and Severe SOR groups. Results confirmed a clear stepwise pattern of increasing internalizing and externalizing symptom burden across the No, Mild, and Severe SOR groups. Specifically, internalizing and externalizing problem scores were significantly greater in the Mild SOR group relative to None (W = 99,562,981 and 90,915,208, respectively; both p < .001). Likewise, internalizing and externalizing problem scores were significantly greater in the Severe SOR group relative to Mild (W = 53,439,657 and 49,334,954, respectively; both p < .001).

**HBN acquisition and preprocessing**

fMRI data were collected with a 64-channel head coil during rest and movie-watching from two locations: CitiGroup Cornell Brain Imaging Center using a Siemens 3T Prisma scanner and Rutgers University Brain Imaging Center using a Siemens 3T Trio scanner. The fMRI data underwent initial minimal processing using the Human Connectome Project minimal processing pipeline.^2^ We conducted further preprocessing utilizing Python 3.8 for both resting-state and movie-watching fMRI data as follows. Raw functional data were denoised using a residual threshold of 0.3 and temporally filtered. To minimize motion-related artifacts, we applied motion correction using framewise displacement (FD), with temporal censoring of volumes exceeding 0.2 mm displacement. The preprocessed data were parcellated according to the Gordon333 atlas and ASEG FreeSurfer subcortical segmentation scheme.^3,4^ For participants with both resting-state and movie-watching scans, data were initially processed separately and later concatenated into a single time series.

**Extending Analysis of FC in ABCD severe, HBN, and ABCD subset matched with HBN**

We additionally tested whether FC-SOR effects for mild SOR in the ABCD exploratory sample 1) generalized to severe SOR among ABCD participants with severe SOR (n=457) and those with no SOR (n=3664) from the ABCD replication group, and 2) generalized to a separate dataset (HBN participants 8-11 years of age, N=356) with distinct sample characteristics. Given sample size constraints, HBN participants were grouped into SOR- (n=175) or SOR+ (n=181) groups, and we conducted a controlled comparison using a matched ABCD subset to account for potential sample size effects. Recognizing that the HBN dataset has a relatively small sample size might limit statistical power to detect effects, we conducted a controlled comparison by creating a matched subset from ABCD with equivalent sample size (n=356), same SOR severity distribution (no SOR=49.2%, mild SOR=36.5%, severe SOR=14.3%), and best matched externalizing symptom profiles (see Fig. S12). Given the relatively small sample size and continuity between effects observed for mild and severe SOR in ABCD, we grouped HBN and ABCD-smaller-subset participants into SOR- (no SOR, n=175) and SOR+ (mild or severe SOR, n=181) groups, rather than analyzing mild and severe SOR separately. Economic disadvantage was operationalized using distinct but comparable measures that were available in ABCD and HBN datasets. This approach allowed us to determine whether differences in FC results between datasets were attributable to sample size limitations rather than true neurobiological differences in SOR-FC associations. Finally, we tested whether the specific SOR-FC associations identified in the ABCD sample extend to the smaller HBN sample mentioned above. Although neither cingulo-opercular-left ventral diencephalon nor somatomotor hand-left hippocampus FC were significantly associated with SOR in the smaller HBN sample (beta=-0.006, p=0.57; beta=0.004, p=0.72, respectively; Table S6), the direction and size of the cingulo-opercular-left ventral diencephalon FC-SOR association was consistent across samples (Figure 5 and Figure S11), according to a Fisher’s r-to-z transformation test (z=0.003).

**SOR-Symptom Analysis by Mild/Severe SOR**

In addition of our main analysis, we further examined the relationship between SOR and psychiatric symptoms across mild-no SOR and severe-no SOR. We similarly subdivided participants into specific age groups to allow for developmental variations, resulting in 8 independent samples and 18 subsamples split by mild/severe (See table S7 for full detail
). For each sample, we conducted separate hierarchical mixed-effects models comparing Mild SOR versus No SOR, and Severe SOR versus No SOR. We separately computed effect-size estimates from conventional bivariate approaches and from our multivariate approach that accounts for common comorbidity/symptom co-occurrence. For bivariate analyses, we tested each psychiatric symptom individually (anxiety, depression, ADHD, ODD, conduct disorder, and autism). In the multivariate analysis, we tested all six psychiatric symptoms together in a single model with SOR. In all models, the dependent variable was binary SOR group (either Mild vs. No SOR, or Severe vs. No SOR), with psychiatric symptoms scores entered as continuous predictors. For datasets that included site information (ABCD, HCP-D, and HBN), we incorporated data collection site as a second-order random-effects variable in the models to account for site-specific variations in analyses.

**Age and sex interaction in SOR*Symptom multivariable models**

To examine whether sex moderated the associations between symptoms and SOR, we in addition included age and sex as fixed variables into our original SOR*Symptoms MLM models to exam if the pattern of result changes and if sex/age show any main effects. When sex and age were added to the primary models, the pattern of symptom associations with SOR remained consistent across all five datasets. We observed no significant main effect in age in any dataset. Regarding sex differences, main effects of sex on SOR were observed in two community samples. Specifically, females had significantly lower odds of SOR compared to males in ABCD (n = 8,854; β = -0.129, p = 0.049) and HCP-D (n = 297; β = -0.84, p = 0.014). In the remaining three datasets (HBN, BGR, SCC), sex did not significantly predict SOR in models accounting for symptom variables and age. We then tested whether the effects of anxiety and autistic traits on SOR differed by sex by including cross-level interactions (predictor × sex). Significant interactions emerged in three datasets: HBN Age 10-11 showed sex moderation for autistic traits, with the association between autistic traits and SOR stronger in males (β = 0.66, 95% CI [0.13, 1.20], p = .017), and for anxiety problems, with the association between anxiety and SOR stronger in females (β = -0.53, 95% CI [-0.99, -0.07], p = .025). HCP-D showed sex moderation for autistic traits, with the association between autistic traits and SOR stronger in males (β = -1.20, 95% CI [-1.98, -0.42], p = .003). ABCD showed sex moderation for anxiety, with the association between anxiety and SOR stronger in males (β = -0.13, 95% CI [-0.26, -0.01], p = .041).

**Cortical and subcortical regions being tested in SOR-functional connectivity analysis**

Segmentations of the following cortical networks and subcortical structures were used to derive cortico-subcortical functional connectivity measures**:** Cingulo-opercular (CON), Sensorimotor hand (SHN), Sensorimotor mouth (SMN), Cingulo-parietal (CPN), Default mode (DMN), Retrosplenial temporal (RST), Salience network (SAL), Visual network (VIS), Dorsal attention (DAN), Fronto-parietal (FPN), Ventral attention (VAN), None network (NN), Ventral diencephalon(VDC), Hippocampus (Hipp), Caudate (Cau), Thalamus(Tha), Nucleus accumbens(Acc), Pallidum(Pall), Putamen(Put), Amygdala (Amyg), Cerebellum (Cereb), Brain stem (Bstem).

**Measures of financial disadvantage in ABCD and HBN used in functional connectivity models**

In the ABCD study, a composite measure was created based on parental responses to seven items assessing eligibility for public assistance programs (e.g., food stamps/SNAP, WIC, free or reduced-price school meals) or inability to afford necessities. Participants were classified as economically disadvantaged if their parents affirmed any of these seven indicators. For the HBN dataset, economic disadvantage was determined using reported annual household income from the Financial Support Questionnaire (FSQ). Households reporting annual incomes below $30,000 were classified as economically disadvantaged, a threshold approximating 150% of the federal poverty level for a family of four during the study period.

**Model selection approach in candidate FC pairs replication analysis**

We identified candidate associations between mild SOR and FC in the exploratory subgroup of the ABCD dataset using hierarchical linear mixed-effects models of FC that accounted for scanner-related variance across multiple study sites as a second-level random-effects variable. We tested for associations with SOR in an exploratory fashion by modeling pairwise (network-network or network-subcortical structure) FC for all FC pairs (n=318). To identify effects that are robust to model selection, each pairwise FC variable was fitted with three separate multilevel models (MLM). For a given FC pair, SOR effects on FC were required to be significant (*p < .01*) and in the same direction for the first 2 models to advance for replication testing. In addition to the second-order variable of scanner, these models contained the following first-order variables:

1. Minimal Model: SOR
2. Primary Model (SOR+Demographics): SOR, demographic variables (Sex, Age, Economic Disadvantage), and mean framewise displacement in retained frames to control for effects of residual motion
3. Symptom Model (SOR+Demographics+Symptoms): SOR, Sex, Age, Economic Disadvantage, Framewise Displacement, and psychiatric measures (internalizing and externalizing behavior scores [CBCL-Internalizing, CBCL-Externalizing] and autistic traits [adjusted S-SRS score excluding SOR])

The FC-SOR associations that replicated in the ABCD study were further tested for extension to FC data from the HBN dataset. The same sequential MLM models were applied to test whether FC-SOR identified in ABCD extended to the HBN sample. We employed a stringent sequential model selection approach to mitigate the multiple comparison issue and potential false positive results FC results due to large sample sizes in ABCD and to systematically evaluate the specificity of SOR-FC associations. Our three-model framework allowed us to test associations at increasing levels of statistical control. The **minimal model** examined whether SOR shows face-valid associations with FC alterations without any covariates. The **primary model** added demographic variables and motion during scan that could largely confound SOR-FC relationships. The **symptom** model further included psychiatric measures (CBCL internalizing/externalizing scores and adjusted SRS autistic traits) to test the specificity of SOR-FC associations beyond general psychopathology.

Given that SOR is highly correlated with clinical symptoms and potentially overlaps with common features across psychiatric conditions, we used the first two models for identifying candidate regions for replication testing. This approach balances the need to control for key confounds while avoiding over-adjustment that could eliminate legitimate SOR-specific effects, particularly important given the substantial comorbidity between SOR and psychiatric symptoms.

**Comparing effect size of FC across samples using R^2^**

R² measures were used to derive effect sizes for FC-SOR effects from the minimal multilevel linear mixed-effects models. The R² (fixed) metric from the r2mlm framework was used to estimate effect size, representing the proportion of total outcome variance explained specifically by SOR via fixed slopes. The R² (fixed) values were calculated using the r2mlm package in R, which implements the integrative framework for multilevel model R² measures described by Shaw et. al .^5^ These effect size estimates allow for direct comparison of FC-SOR effects from the Minimal Model across FC pairs and between the ABCD and HBN cohorts.

**Confirming that replicating FC-SOR effects are not due to family-level clustering**

To ensure that the FC-SOR results are not inflated due to within-family dependencies between the subsamples, we repeated the analysis with an alternative matched ABCD subsamples stratified at the family level to enforce maximal independence between exploratory (ARMS-1) and replication (ARMS-2) samples. This approach built upon the ARMS methodology^6^ which accounts for family structure during demographic matching, but extended it to enforce complete family-level independence for all statistical comparisons and permutation testing. Specifically, all siblings and family members were assigned as a unit to a single subset, preventing families from being divided across the exploratory and replication samples.

*Model Results for CPN-right and left Caudate*

Using family-level stratified ARMS subsets, we examined whether the cingulo-parietal network to bilateral caudate findings replicated while enforcing complete family-level independence. These two key connections showed the same robust replication across exploratory (AsRMS-1) and replication (ARMS-2) subsamples, consistent with the results we found in our original subsamples (ABCD exploratory and ABCD replication): **CPN-right caudate FC was** significantly negatively associated with mild SOR in both ARMS-1 (β = -0.014 to -0.017, p < 0.01 **across all 3 models**) and ARMS-2 (β = -0.020, **p < .0001 across all 3 models**). Similarly, **CPN-left caudate FC was significantly negatively associated with mild SOR in both** ARMS-1 (β = -0.012 to -0.015, p <0.05) and ARMS-2 sample (β = -0.023 to -0.025, **p < .00002**), indicating that the observed replicating effects in the main analyses are not attributable to family-level data leakage across subsamples.

*Correlations of brain-wide FC-SOR effects are significantly higher than permuted correlations for ABCD-ARMS1-to-ABCD-ARMS2 and ABCD-ARMS1-to-ABCD-Severe samples, but not ABCD-ARMS1-to-HBN*

As in the main analyses, the mean of the 1000 permuted ARMS1-to-ARMS2 correlations and ARMS1-to-Severe correlations are significantly lower than the actual correlations (ARMS1-ARMS2 actual rho=0.36, Permuted rho=-0.09, p=0.012; ARMS1-Severe actual rho=0.53, Permuted rho=-0.010, p<0.0001). As in the main analyses, the actual ARMS1-to-HBN Spearman’s correlation is not significantly different from the permuted correlations (actual rho=-0.016, Permuted rho=-0.0017, p=0.5, see figure S10).

**Supplementary Tables and Figures**

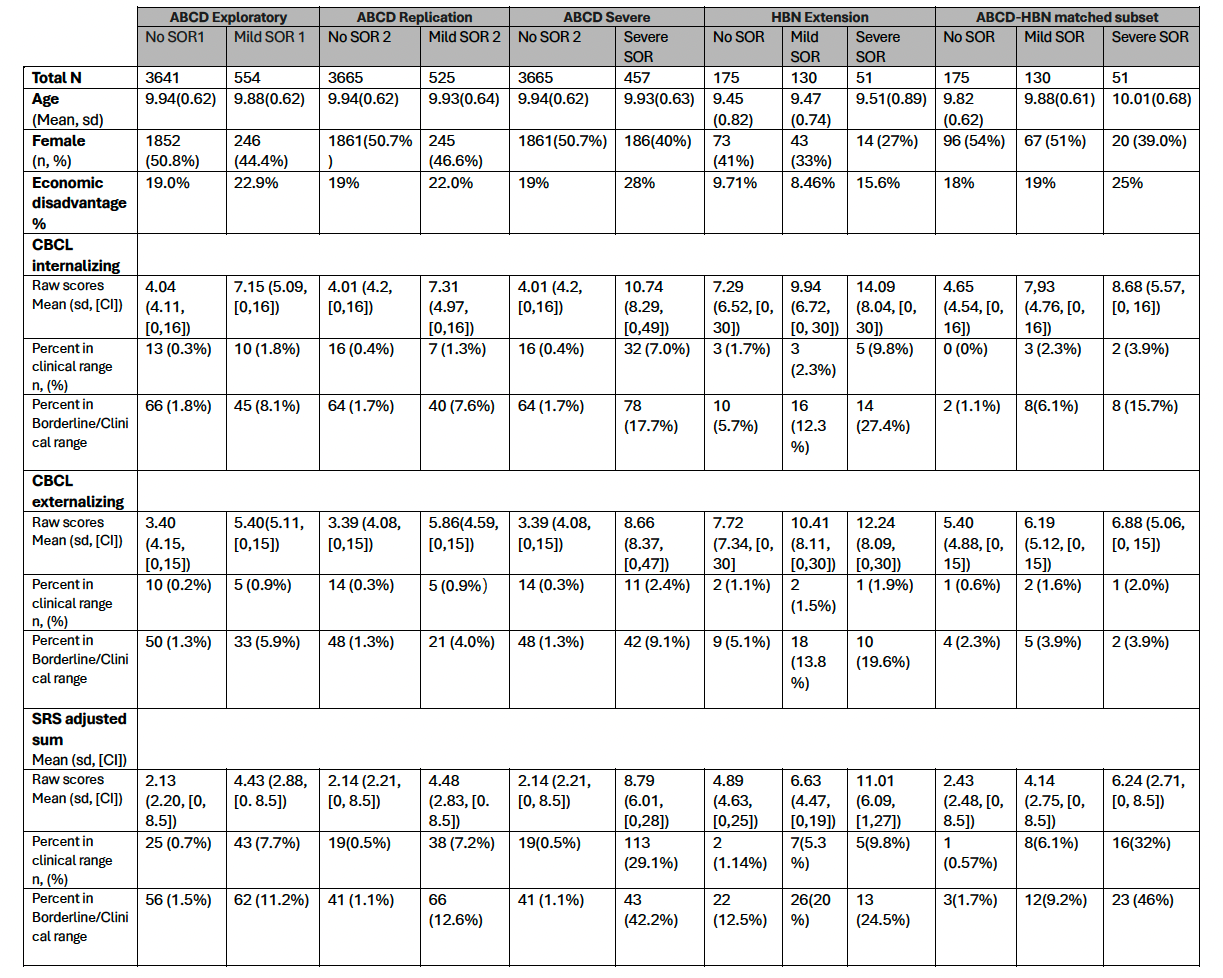
**Table S1:** Demographic information for subsamples in functional connectivity analyses.

Percentages of children with clinically significant baseline psychiatric symptom scores in the full

sample and in each SOR group are shown. CBCL T-scores for internalizing and externalizing, S-SRS/SRS (based on data availabilities across datasets) T-scores for autism spectrum disorders were used to identify clinical significance with scores in the clinical range defined as T≥70 and scores in the borderline clinical or clinical range defined as T≥65.

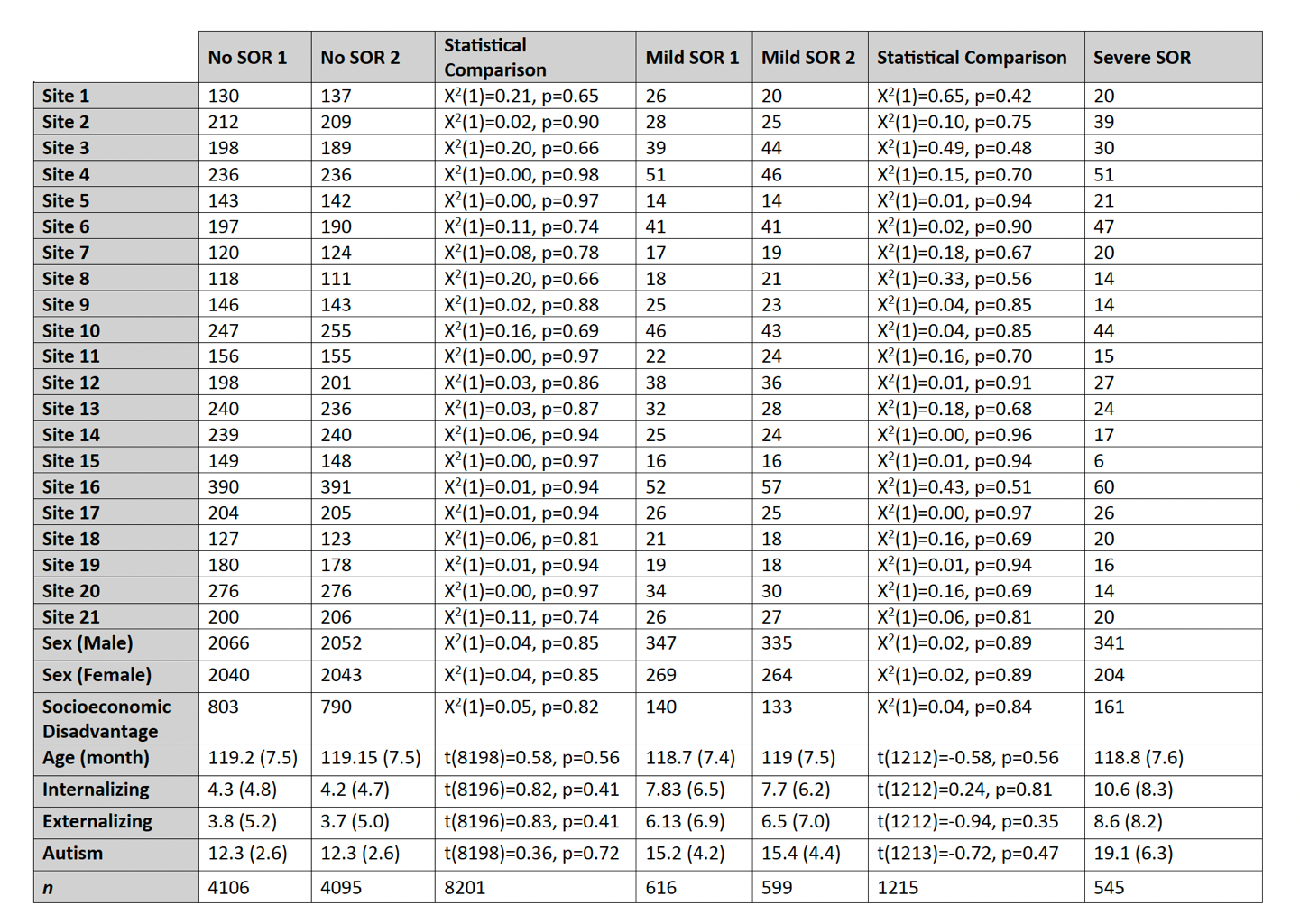
**Table S2:** The participants from ABCD Exploratory and Replication from are matched in sample sizes, SOR prevalence, sex, socioeconomic disadvantage, age, clinical symptoms, and autistic traits by study site.

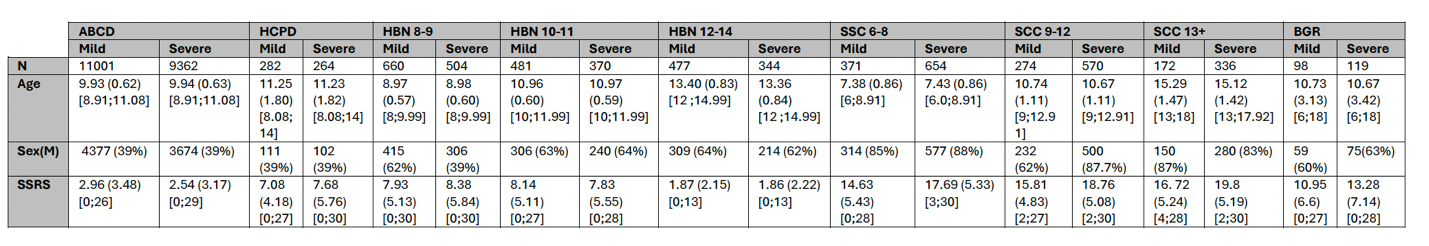
**Table S3:** Age, Sex, and SRS scores of nine subsets of participants derived from the five study samples. Participants from two datasets (i.e., HBN, SSC) with larger age ranges and sample sizes were divided into subgroups by age.

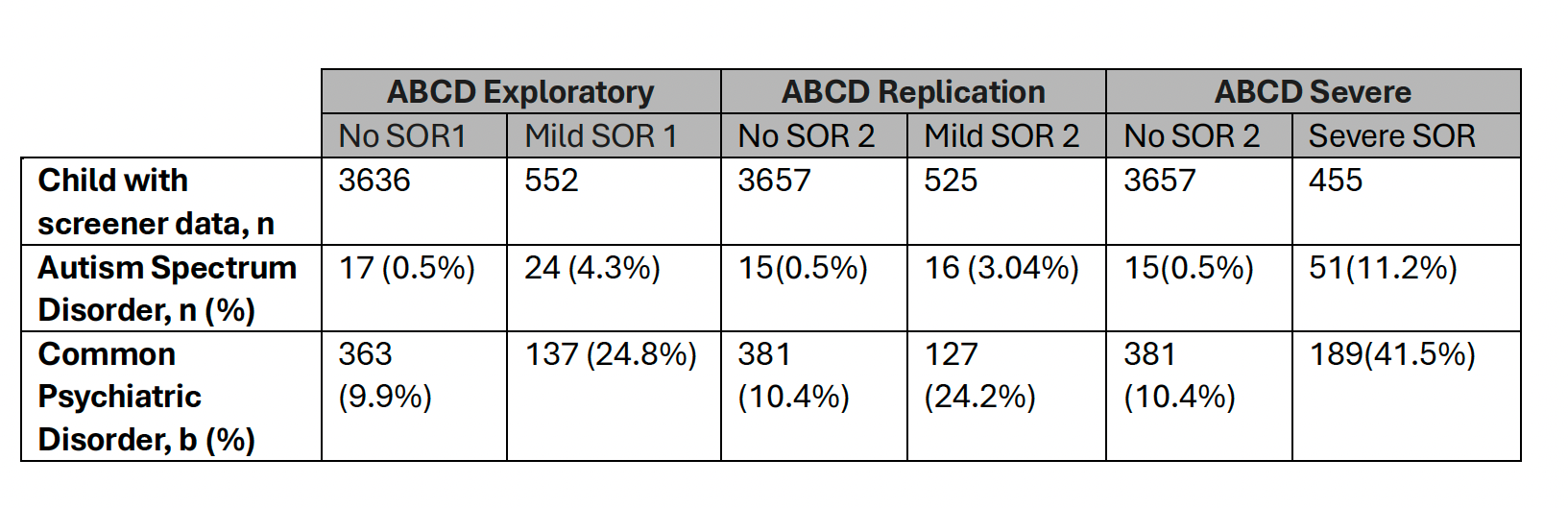

**Table S4:** Parent report of child diagnoses on screener in ABCD subjects used in FC-SOR analysis broken down by no/mild/severe SOR. Parents completed a screener prior to child participation in the ABCD study. Screener questions included whether the child has ever received a diagnosis of autism spectrum disorder and whether the child has ever received a diagnosis of a psychiatric disorder like ADHD, depression, bipolar disorder, anxiety disorder, or phobia. This table shows the number and percent of children in each SOR group (no/mild/severe) from the FC analysis who had received such diagnoses by the screener (pre-Y0) according to parent report. Note that sample sizes for children with screener data could be smaller than sample sizes for analyses, reflecting the fact that screener responses were not available for all participants.

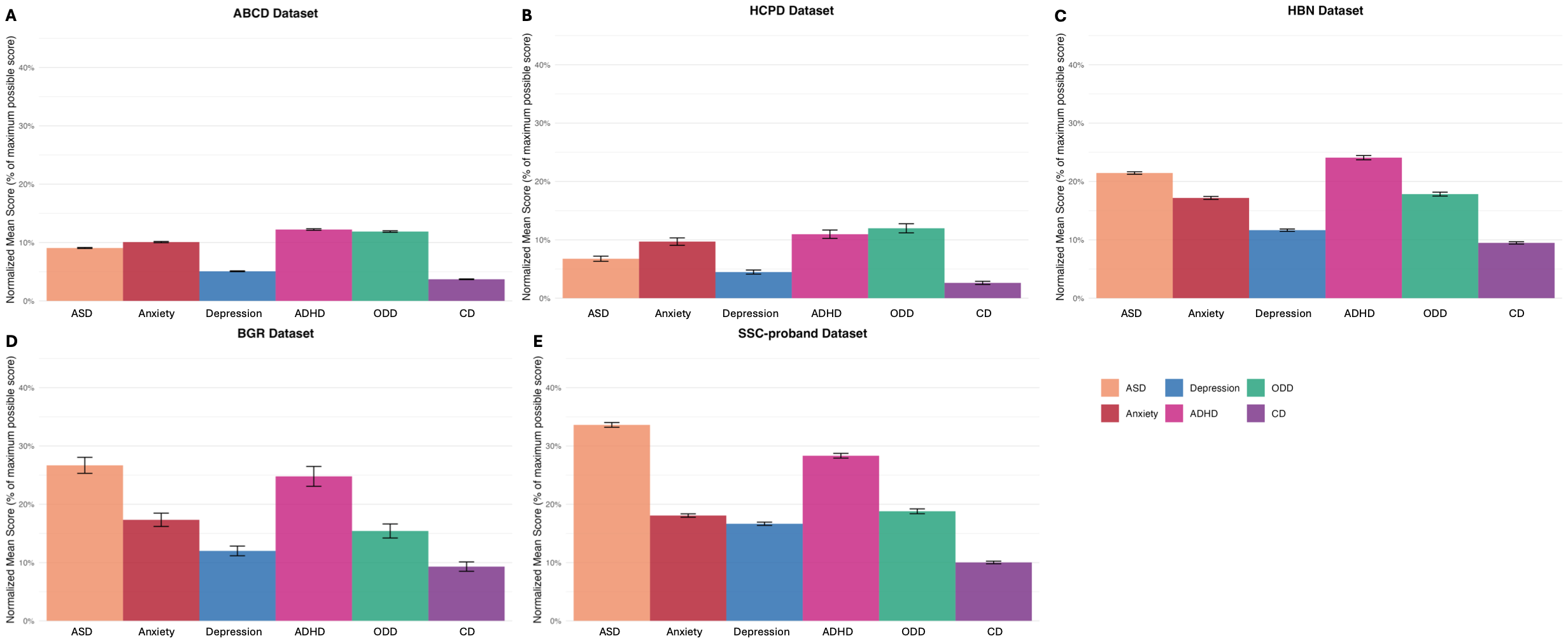

**Figure S1: The psychiatric symptom profile across five datasets is distinct from each other especially between community samples (ABCD, HCP-D, HBN) and ASD samples (BGR, SCC): A.** Normalized mean scores of ASD, anxiety, depression, CD, ODD in ABCD dataset. **B.** Normalized mean scores of ASD, anxiety, depression, CD, ODD in HCP-D dataset. **C.** Normalized mean scores of ASD, anxiety, depression, CD, ODD in HBN dataset. **D.** Normalized mean scores of ASD, anxiety, depression, CD, ODD in BGR dataset. **E.** Normalized mean scores of ASD, anxiety, depression, CD, ODD in SCC dataset.

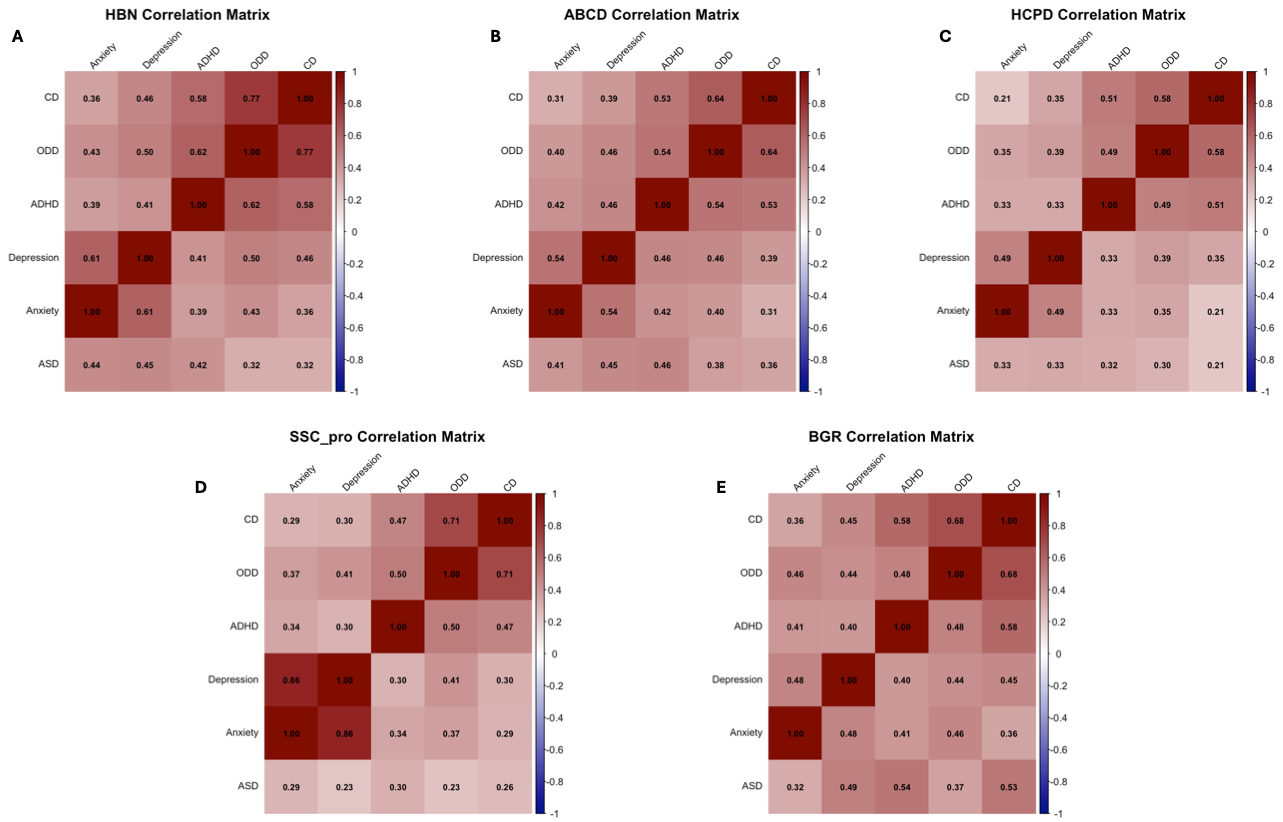

**Figure S2: All the psychiatric symptoms are positively correlated with each other within each dataset:** **A.** Spearman correlation matrix between psychiatric symptoms for HBN study. **B.** Spearman correlation matrix between psychiatric symptoms in ABCD study. **C.** Spearman correlation matrix between psychiatric symptoms in HCPD study. **D.** Spearman correlation matrix between psychiatric symptoms in SCC study. **E.** Spearman correlation matrix between psychiatric symptoms in BGR study.

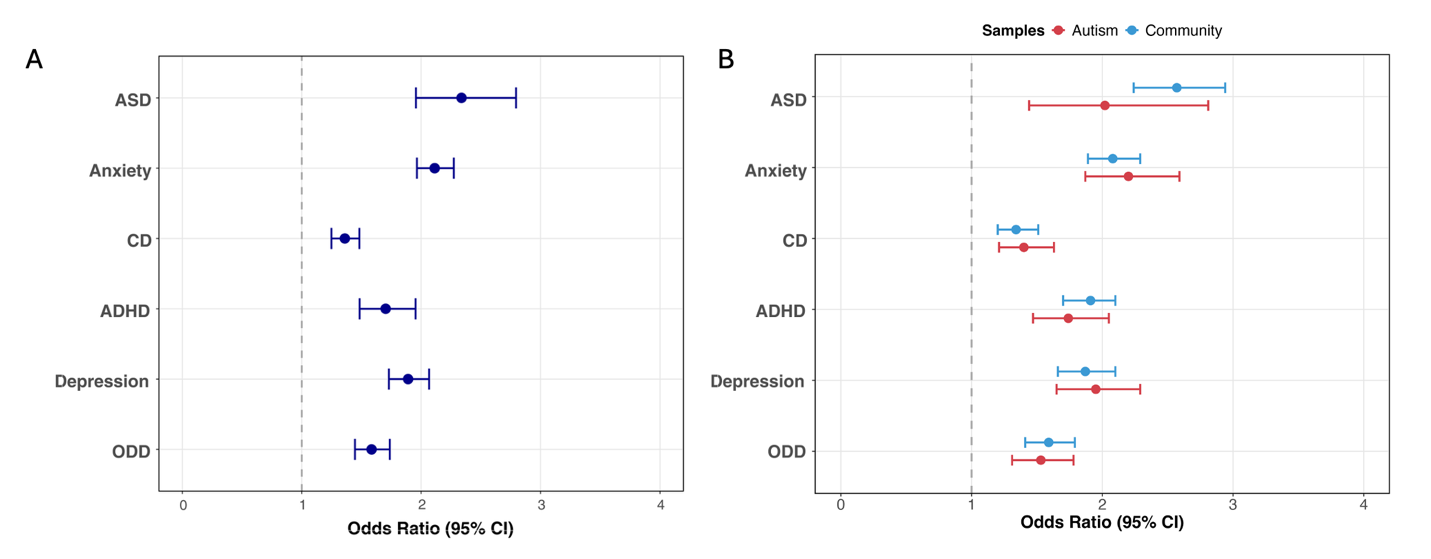

**Figure S3: SOR is significantly positivity associated with all the psychiatric symptoms from the bivariate models across five datasets. A.** Bivariate associations between SOR and psychiatric symptoms pooled across all datasets. **B.** Bivariate associations between SOR and psychiatric symptoms pooled by community (ABCD, HBN, HCP-D) and autism-enriched (BGR, SSC) datasets.

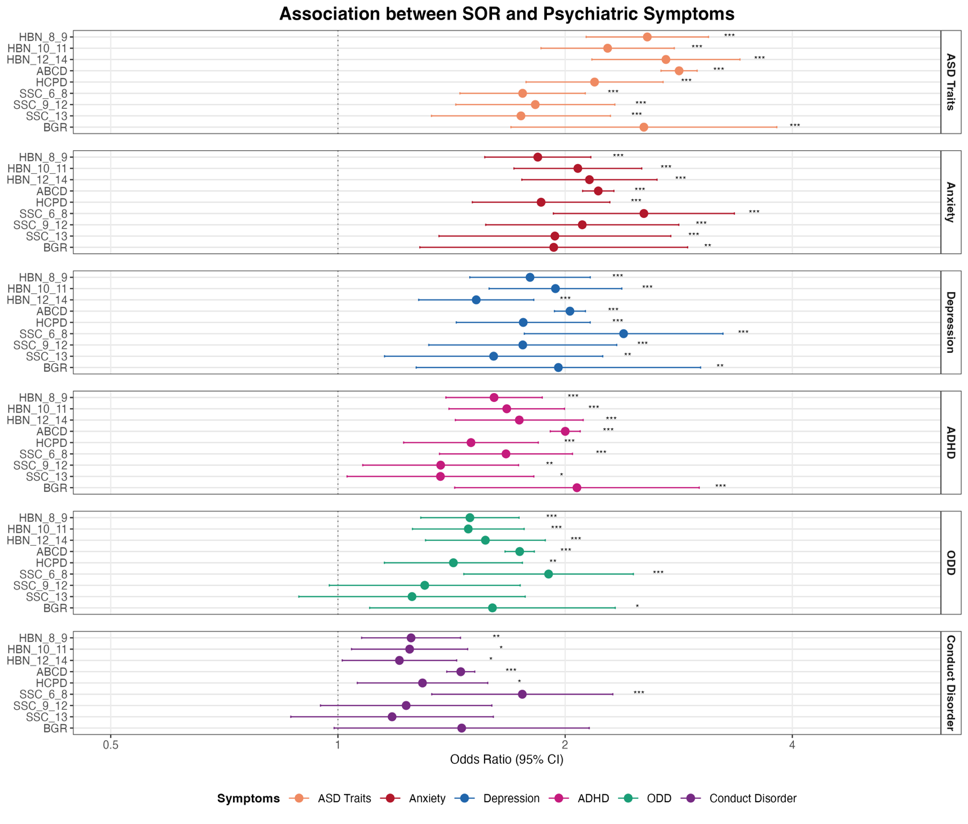

**Figure S4:** SOR is positively correlated with ASD, anxiety, depression and ADHD across all 9 subsamples different in age and datasets in bivariate models. It is also positively associated with ODD and CD in all subsamples besides older subjects from SCC and BGR.

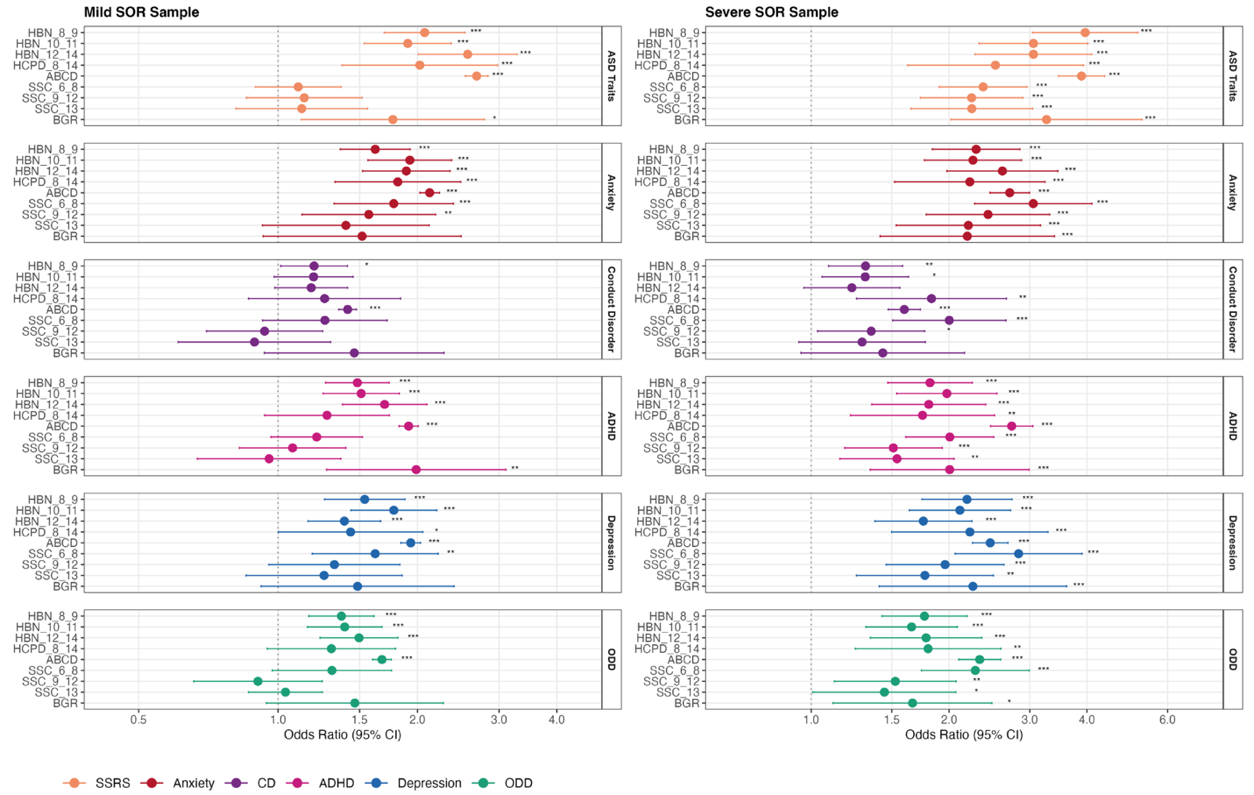

**Figure S5:** Specific odds ratios from the bivariate symptom*SOR models in all subsamples by mild-no and severe-no SOR shows that SOR is positively associated with all six neurodevelopmental traits and psychiatric symptoms in most samples regardless of SOR severity.

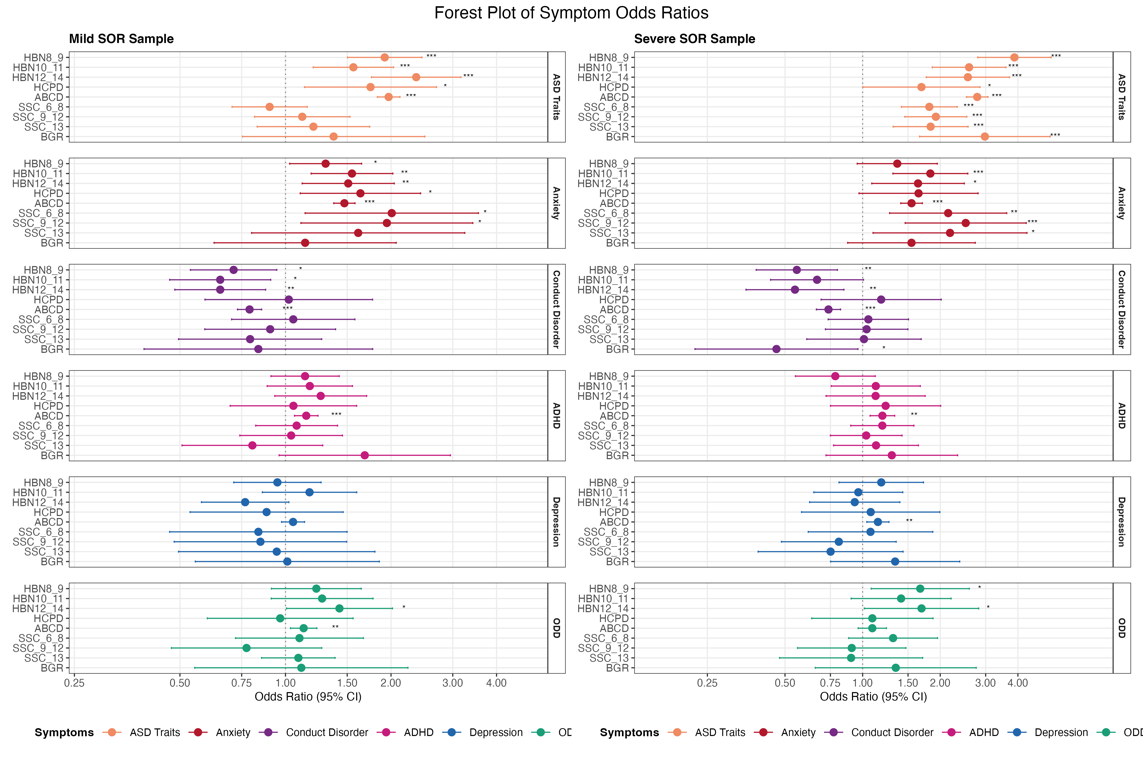

**Figure S6:** Specific odds ratios from the bivariate symptom*SOR models in all subsamples by mild-no and severe-no SOR shows that SOR is only consistently positively associated with autistic traits and anxiety problems regardless of SOR severity and negatively associated with conduct disorder problems in most community samples.

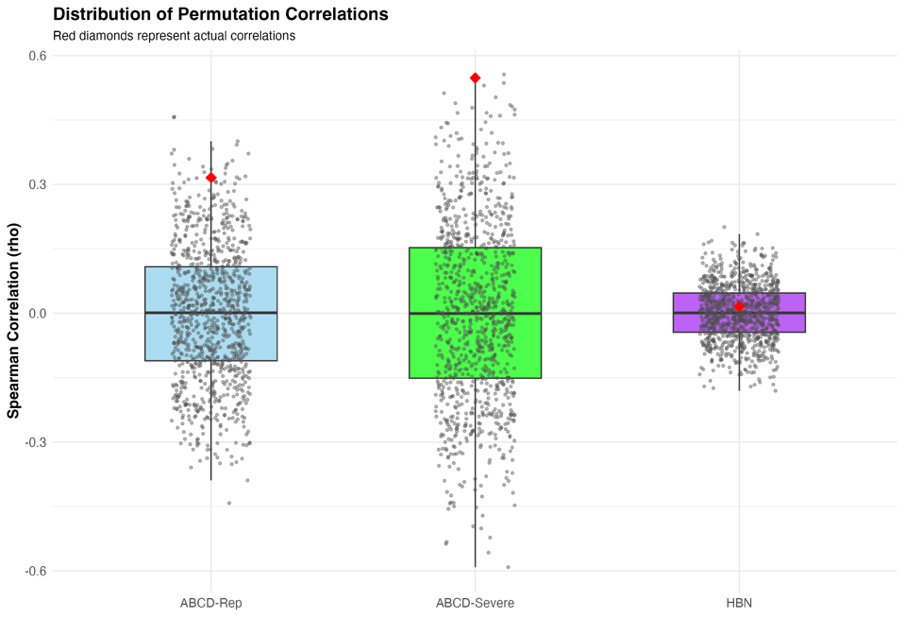

**Figure S7: The actual correlations of brain-wide FC-SOR effects are significantly higher than permuted correlations for ABCD exploratory-to-replication and ABCD exploratory-to-severe, but not ABCD exploratory-to-HBN:** Distribution of Spearman correlations of the ABCD-exploratory beta matrix with 1000 permuted versions of ABCD-replication (left), ABCD-severe (middle), and HBN (right) beta matrices are shown with black dots. Spearman correlations of ABCD-exploratory beta matrix with actual ABCD-replication, ABCD-severe, and HBN beta matrices are shown with red diamonds.

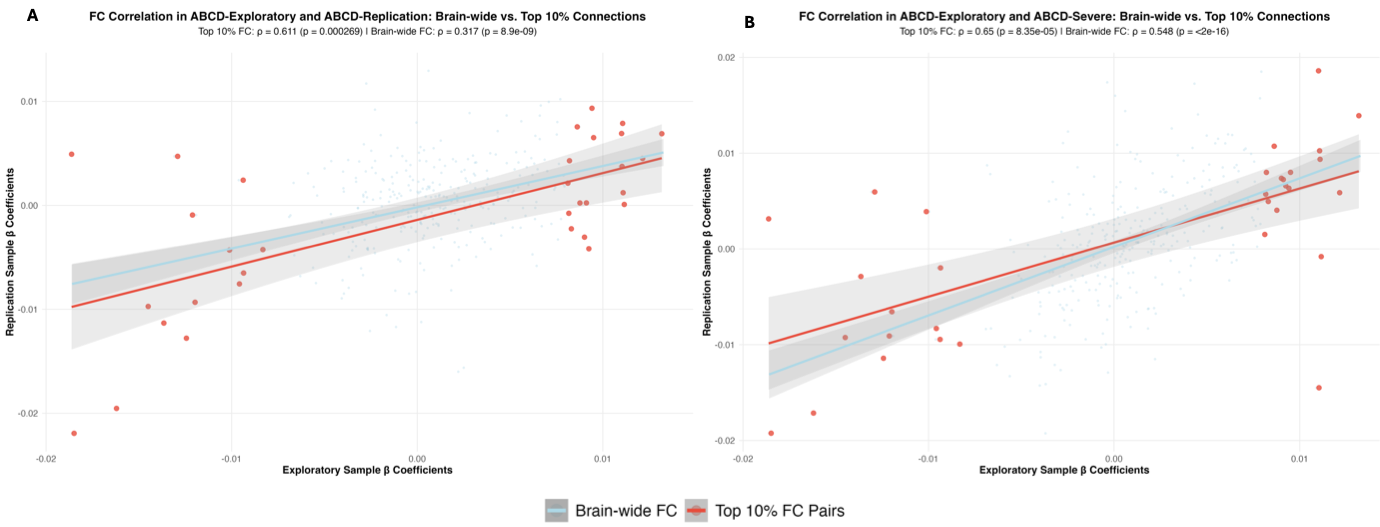

**Figure S8:** **Correlation coefficients for the largest 10% FC-SOR effects from the ABCD-Exploratory Beta Matrix: Relation to ABCD-Replication and ABCD-Severe effects:** A. Spearman correlations of the top 10% of largest FC-SOR effects versus the brain-wide FC pattern associated with SOR between ABCD exploratory-ABCD replication subsets (ρ = 0.61, p < 0.001). B. Spearman correlations of the top 10% of largest FC-SOR effects versus the brain wide functional connectivity pattern associated with SOR between ABCD exploratory-ABCD Severe subsets (ρ = 0.65, p < 0.001).

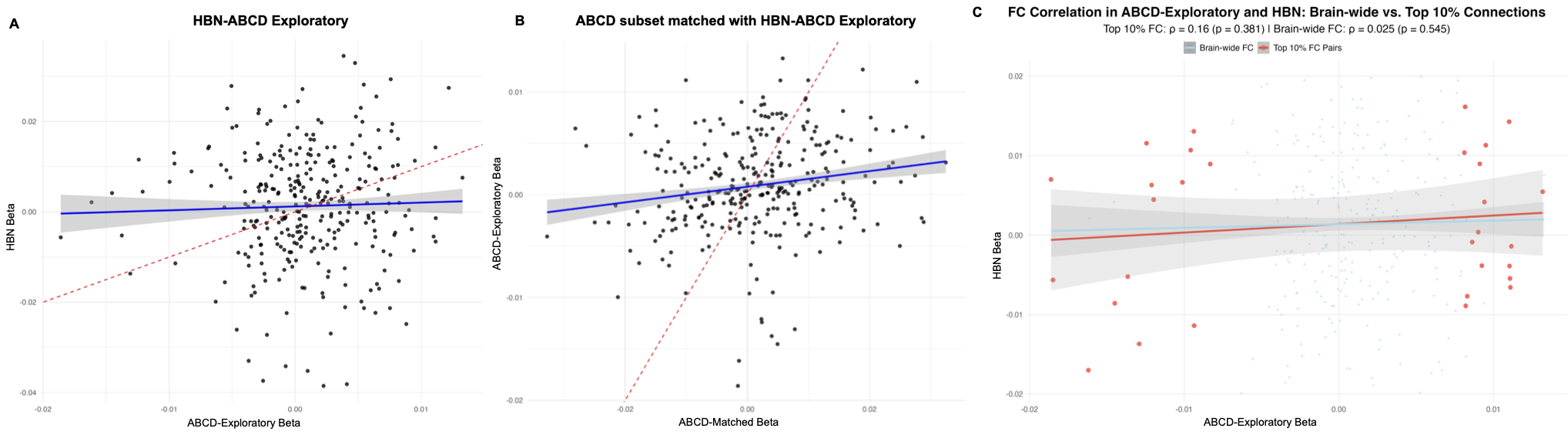

**Figure S9: Brain-wide FC pattern found in ABCD-exploratory does not extended to HBN dataset or a subset of ABCD sample matched to HBN: A.** Spearman correlation between ABCD-Exploratory and HBN beta matrix (rho=0.025, p=0.545). **B.** Spearman correlation between the subset of ABCD-matched HBN sample beta matrix and ABCD-Exploratory (rho=0.174, p=0.07). C. Spearman correlations of the top 10% of largest FC-SOR effects versus the brain-wide FC pattern associated with SOR between ABCD exploratory- HBN (rho=0.16, p=0.381). None of the candidate FC pairs we have identified in ABCD-Exploratory were significant in this subset as well. These results suggest that larger sample sizes are needed to derive replicable estimates of brain-wide FC patterns associated with SOR.

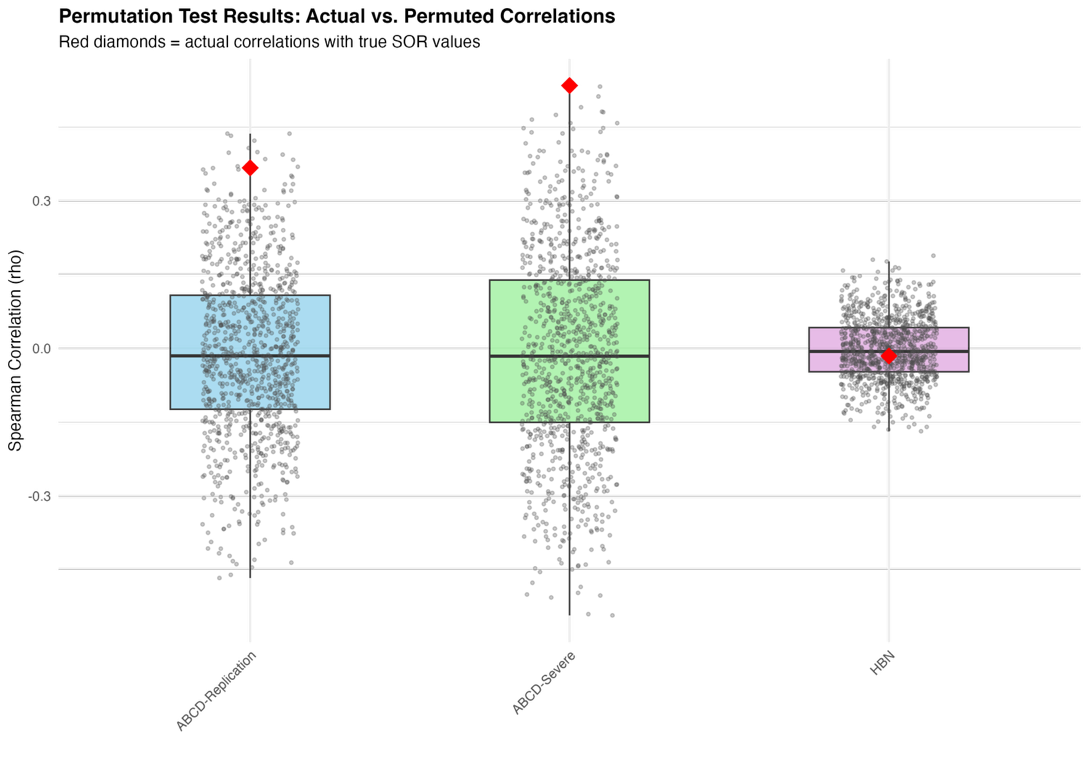

**Figure S10: The correlation coefficients between the whole-brain FC-SOR effects and top 10% of largest FC-SOR effects from the ABCD-ARM1 to ABCD-ARM2, ABCD-Severe, and HBN**

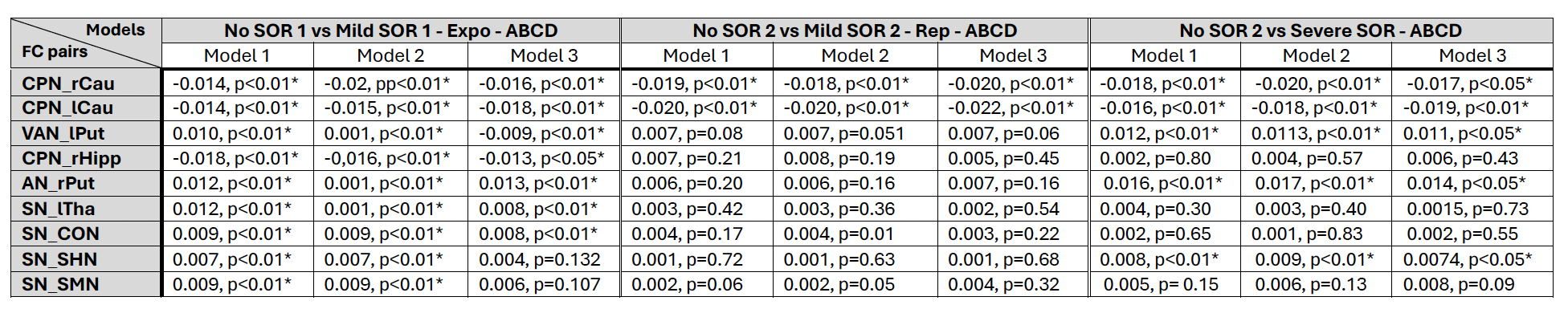

**Table S5:** FC Model results of the 9 candidate FC pair identified in ABCD-exploratory from all ABCD subgroups

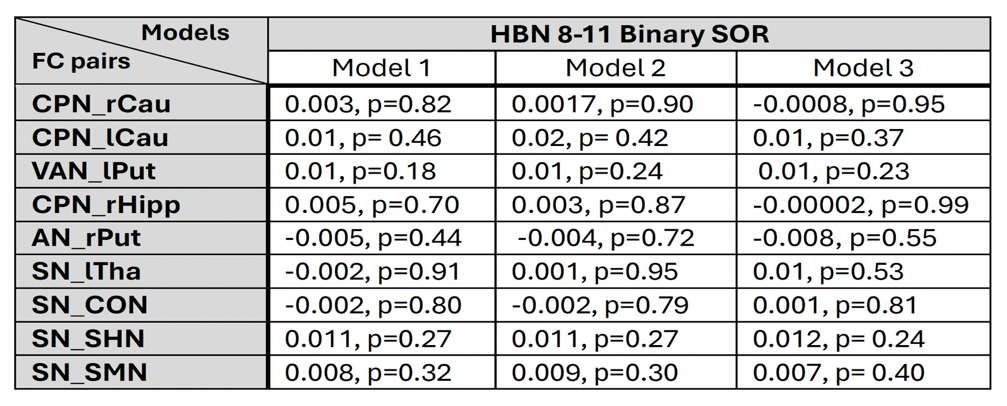

**Table S6:** FC model results for HBN of the 9 candidate FC pair identified in ABCD-exploratory

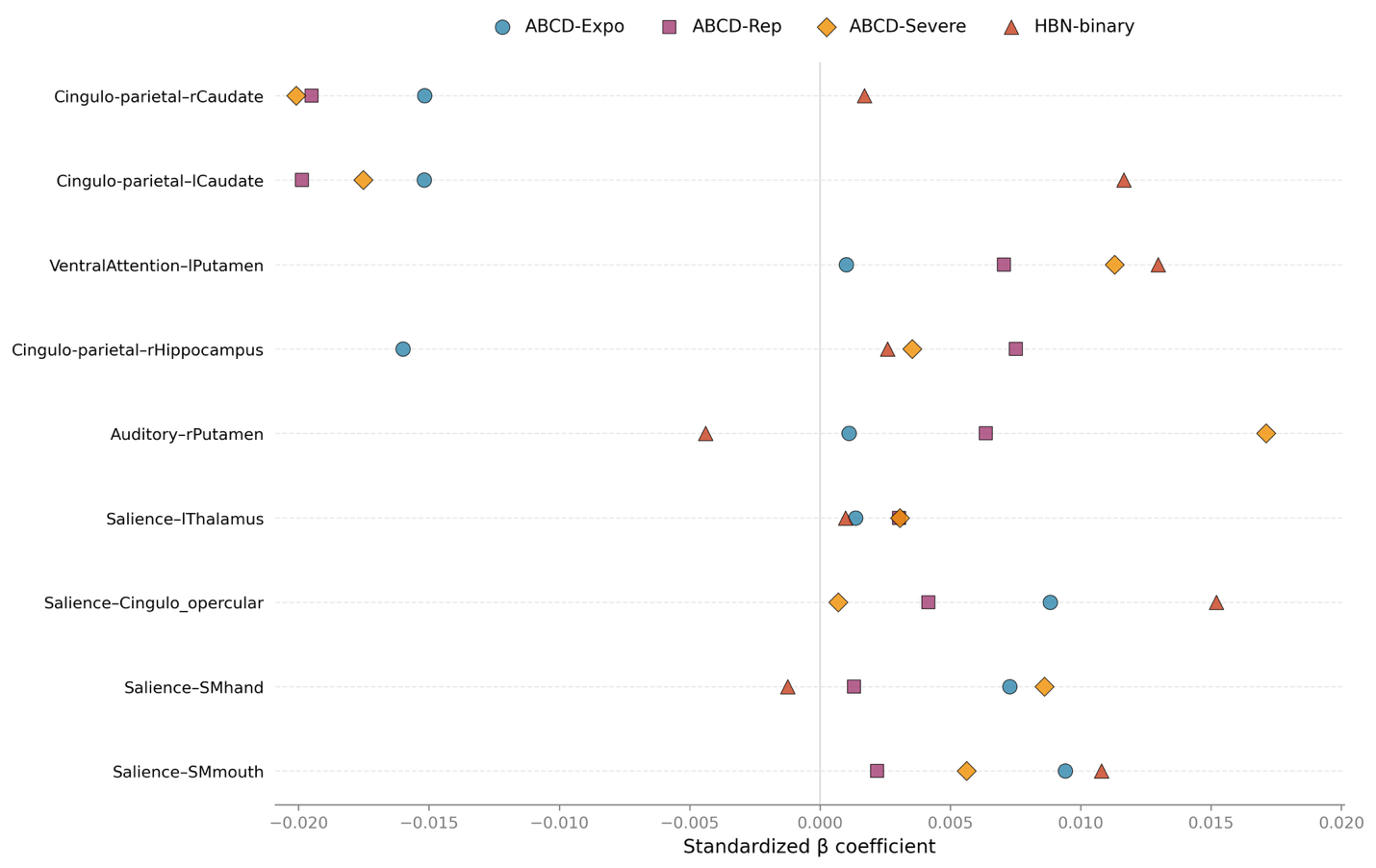

**Figure S11: Summary of Fixed Effect R² Values Across ABCD**: Fixed effect R² values for the nine functional connectivity pairs tested across three samples: ABCD-exploratory, ABCD-replication, ABCD-severe.

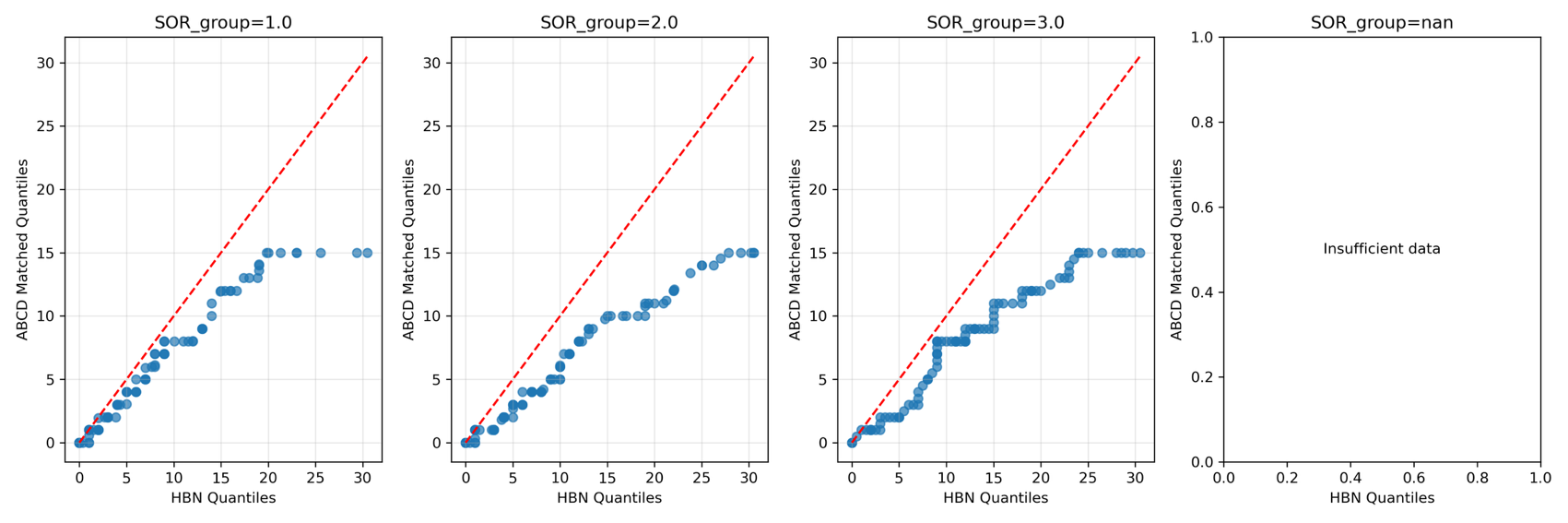

**Figure S12**: **The HBN and ABCD-matched HBN subsample are highly similar in externalizing symptoms scores distribution across no SOR group (SOR_group=1.0), mild SOR group (SOR group=2.0), and severe SOR group (SOR group=3.0):** Quantile-quantile plots comparing ABCD and HBN CBCL-externalizing raw symptom scores distributions across SOR severity groups.
